# Magnetically Controlled Microrobots for In Vivo Non-Invasive Embryo Transfer

**DOI:** 10.1101/2025.09.29.679217

**Authors:** Azaam Aziz, David Castellanos-Robles, Zhi Chen, Ronald Naumann, Harini Raghu Kumar, Richard Nauber, Carla Ribeiro, Ripla Arora, Mariana Medina Sánchez

## Abstract

Infertility affects millions worldwide and is often linked to factors such as poor sperm quality and female reproductive organ disorders. Despite significant advancements in in vitro fertilization (IVF) and intracytoplasmic sperm injection (ICSI), implantation rates remain low, ranging from 17 to 21% after three days of incubation, mainly due to stress, lifestyle factors, and uterine conditions. Extended embryo culture techniques have shown promise in improving pregnancy rates. However, the availability of high-quality blastocysts remains a major challenge. Intrafallopian transfer techniques, such as gamete/zygote intrafallopian transfer (GIFT/ZIFT), were introduced to improve fertilization and early embryo development, particularly for patients with repeated embryo implantation failure (10–30% of assisted reproduction technology (ART) cases, particularly in women >35). However, these methods have declined due to advancements in IVF and the variability in laparoscopy procedures used for GIFT/ZIFT. To address these challenges, we propose a non-invasive microrobotic embryo transfer (µET) technique using remotely controlled microcarriers comparable in size to embryos. We demonstrate the capabilities of magnetically actuated spiral microrobots, fabricated using laser direct writing for capturing, transporting, and releasing embryos into murine uteri. Additionally, we characterize their motion performance and implement image-guided closed-loop control and dual ultrasound (US)/photoacoustic (PA) tracking for deep-tissue interventions. Recognizing the importance of clinical translation, we present preliminary studies on gelatin-based microrobots, a biodegradable alternative, and explore endometrial remodeling after the in vivo transfer of microrobots carrying embryo-like structures. Our findings show that these microrobots can effectively transport embryos, support their development, and enable minimally invasive delivery, providing a more natural, targeted, and non-invasive strategy for in vivo assisted reproduction.

**One-Sentence Summary:** We propose a non-invasive µET technique using magnetically actuated microcarriers to improve embryo cargo-delivery in assisted reproduction, addressing challenges in infertility treatments by enabling precise, image-guided embryo delivery with minimal invasiveness.

## INTRODUCTION

Infertility affects millions of couples worldwide^1^, stemming from various causes, including poor sperm quality, ovulatory disorders, and uterine abnormalities. Despite improvements in IVF and ICSI reaching fertilization rates of about 95%^2^, embryo implantation remains a key bottleneck with rates from 17 to 21%^3^, especially with advancing age.^4^ Extended embryo cultivation has led to higher pregnancy rates of 42 to 47%^4^ but obtaining high-quality blastocysts remains challenging, necessitating numerous oocytes and advanced assessment techniques. Low pregnancy rates may stem from oxidative stress affecting gametes during in vitro manipulation, as well as from patient lifestyle factors, underlying diseases, or uterine abnormalities. GIFT/ZIFT offers a more physiological environment for oocyte fertilization and embryo development. It allows for the transfer of gametes or early embryos back to the fallopian tube via laparoscopy, showing potential benefits for recurrent embryo implantation failure (RIF), a problem affecting 10 to 30% of couples undergoing fertility treatments^5^. However, laparoscopy requires centimeter-scale incisions and multiple instruments, carrying risks of ectopic pregnancy, infection, and surgical injury, and outcomes remain variable, depending on surgical expertise and laboratory protocols. This underscores the need for less invasive, more precise approaches for embryo transfer (ET).

Recent advances in medical technology, including miniaturized devices such as catheters and microrobots, now enable the development of cellular-scale tools that can be remotely controlled using safe physical fields like magnetic or acoustic waves, allowing for minimally or non-invasive navigation through complex and hard-to-reach environments within the body^6,7^. Microrobotic carriers on the cellular size scale could therefore address RIF by transporting gametes, early embryos, and therapeutic cargoes to the ampulla site of the fallopian tube, supporting embryo development in a more natural setting while reducing the process invasiveness. Similar microrobots have been implemented in various other medical applications such as targeted drug delivery^8–11^, minimally invasive surgery^12^, blood clot removal^13^, or cell transport^14^ among others, even in living organisms. Microrobots are actuated and guided by external physical means i.e., magnetic fields, ultrasound, or light for precise remote manipulation and tracking to perform the intended functional tasks.

Current assisted reproduction methods lack the ability for accurate ET and rely heavily on the doctor’s expertise and the patient’s tolerance for invasive procedures. Given that the success rate of ICSI exceeds 90%, while the rate of successful pregnancies following ET is just over 30%, it is evident that embryonic development after fertilization and the ET procedure are key determinants of overall success^15^. Therefore, microrobotic tools and non-invasive methods for gamete and embryo manipulation hold great potential for improving pregnancy rates. Crucially, these microrobots must effectively capture and securely transport gametes or embryos while allowing access to secreted molecules, growth factors, antioxidants, from the reproductive tract. Unlike conventional ET protocols in ART, which primarily rely on transcervical catheter-based techniques, these methods have inherent limitations and invasive aspects. Standard ET involves inserting a catheter through the cervix into the uterus to deposit the embryo, a procedure that can cause discomfort, cervical trauma, undesired immunoreactions, or uterine contractions, potentially lowering implantation success rates. Additionally, factors such as catheter rigidity, operator expertise, and endometrial receptivity play a crucial role in determining the pregnancy outcome.

µET offers a less invasive alternative for delivering embryos to the reproductive tract, either to the uterine cavity (blastocysts) or the fallopian tubes (zygotes). For oncological patients who are unable to undergo hormonal stimulation, transporting embryos to the reproductive tract during the natural cycle could offer significant benefits. Our group has previously reported a preliminary work on capturing zygotes using microrobots^16^. This proof-of-concept study successfully demonstrated the transport and release of a single murine embryo within microfluidic channels, showcasing the carriers’ ability to securely retain zygotes during the transition from the Petri dish to the cannula, critical steps for subsequent intrauterine or intrafallopian delivery. However, several challenges needed to be addressed, including validating the functionality of these microcarriers in living models, such as mice, and demonstrating their capacity to transport multiple embryos. This is particularly important, as murine ET typically requires at least nine embryos for optimal pregnancy success. In human applications, the ultimate goal is to enable single ET to reduce the risk of multiple pregnancies.

To validate the potential of the µET technique presented here, it is essential to visualize the ET process within the uterus. Medical imaging or tracking of such microrobots is not straightforward in the complex in vivo environment. It poses significant limitations in achieving high spatiotemporal resolution and precise anatomical positioning within living organisms. There have been several attempts using various imaging techniques to monitor microrobots in vivo^17–25^. However, to address the need for an imaging system capable of real-time monitoring and non-invasive observation over an extended period, we adopted a dual US/PA imaging approach. This choice is particularly pertinent as our microrobots, with a size of approx. 300 µm differs from tissue and body fluids in that they are coated with plasmonic materials, resulting in a strong PA signal for enhanced tracking capabilities.

In this study, we demonstrate for the first time a complete µET workflow using magnetically actuated microrobots for capturing, transporting, and releasing blastocysts into mouse uteri. It highlights their efficient propulsion across tissue surfaces and within confined channels, as well as their closed-loop control using both optical and PA imaging. The spirals are able to securely hold one or multiple embryos, effectively creating a microenvironment around the embryo during transport while still allowing exposure to the surrounding medium (so the embryo can benefit from uterine secretions). We demonstrate that embryos can be successfully loaded into microrobots and cultured in vitro within them, progressing from the 1-cell stage to the blastocyst stage (prior to transfer). Furthermore, we present a complete workflow, from in vitro loading of pseudo-embryos into the spiral microrobots, to their intrauterine transfer using a cannula, followed by external magnetic guidance. Histological analysis and confocal microscopy confirmed the apparent biocompatibility of the materials and structures, showing no evident toxic effects on surrounding tissues. These analyses also reveal uterine tissue remodeling and lining around the released microrobot in pseudopregnant mice.

Furthermore, recognizing the need for materials that are both biocompatible and biodegradable, aligned with the ultimate goal of the proposed µET approach—we explored the use of gelatin for fabricating the same microrobot designs. We demonstrate that these microrobots can support embryo development while degrading and assessed their potential for future clinical translation.

In summary, our µET method introduces a disruptive approach to assisted reproduction by enabling precise, minimally invasive embryo transport with magnetically actuated, cellular scale microrobots. Combined with real-time, image-guided control, this approach has the potential to enhance ET precision, improve retention and implantation, and increase safety, paving the way for the next generation of in vivo assisted reproduction techniques.

## RESULTS

### Microrobotic ET Concept

The concept of minimally invasive, microrobotic ET in vivo is illustrated in **Fig. 1a**. A cannula carrying a microrobotic payload is inserted through the vaginal opening into the uterine cavity. Once the embryo is guided by external rotating fields to deeper locations within the uterine horns and released from the spiral-shaped microrobot, it can continue developing under natural conditions. The transfer procedure utilizes a spiral microrobot capable of capturing, transporting, and releasing one or multiple embryos. To demonstrate the feasibility of the concept, a mouse model was chosen, as multiple pregnancies can be achieved by delivering approx. 6–9 embryos to the uterus. In contrast, human ET typically involves a single embryo. The procedure is performed under rotating magnetic control and is fully reversible. To achieve this, a spiral-shaped microrobot (∼300 µm footprint, 110 µm height) was designed to enclose and protect up to three embryos while maintaining contact with the surrounding fluid. The goal is to transport 2–3 such spirals, each carrying three embryos, to both uterine horns. The microrobot’s motion mechanism enables precise embryo capture at its center and controlled release by reversing its rotational direction. The spirals are fabricated using two-photon absorption lithography and designed to fit mouse zygotes/embryos (∼60–90 µm). They must be large enough to enclose the embryos securely but small enough to navigate the uterine horns. The spiral’s opening is designed to guide the embryo toward the center while preventing escape. The optimal spiral microrobot design was achieved after several iterative refinements as depicted in **Fig. S1.**

**Fig. 1.**
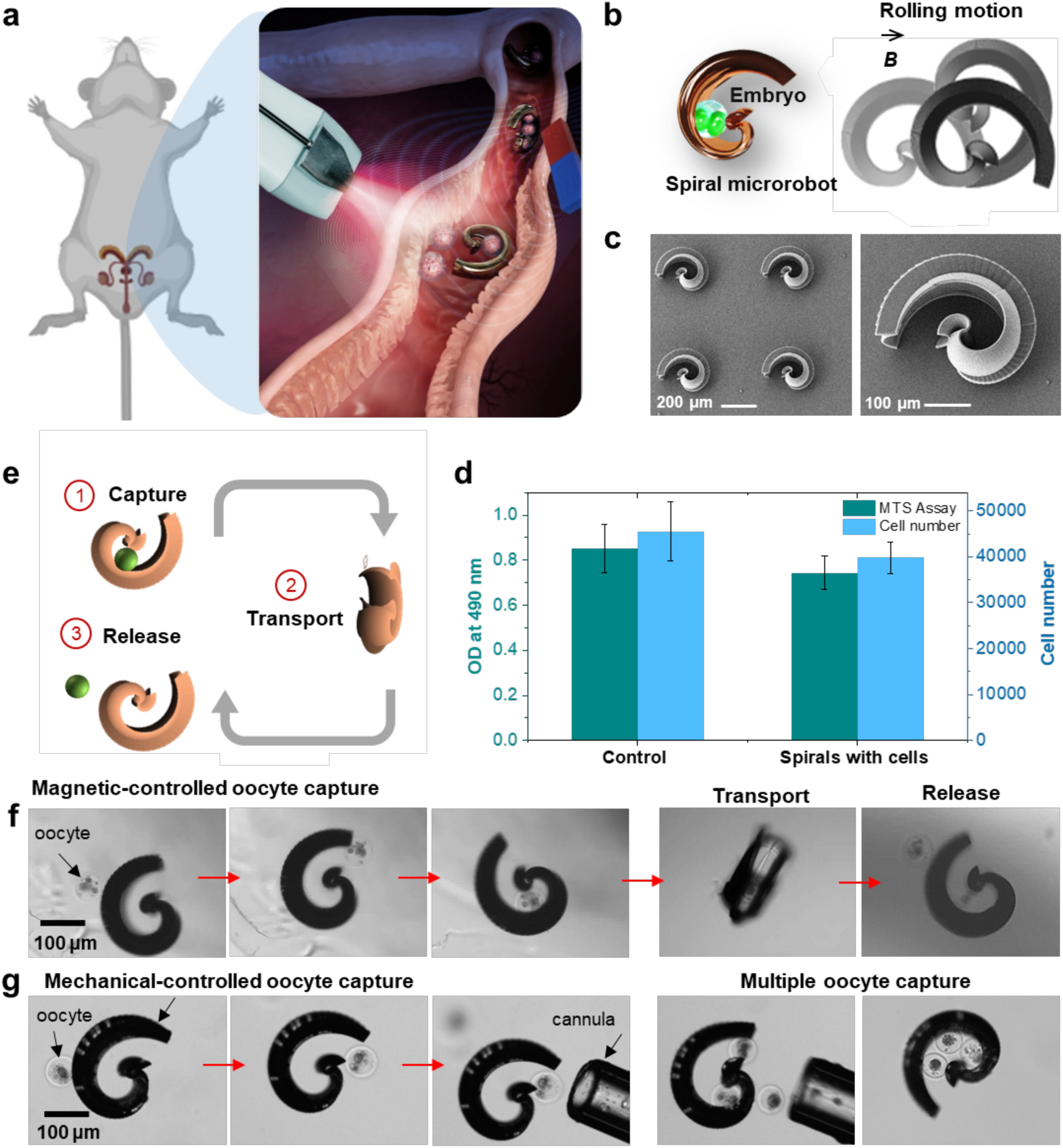
Microrobotic embryo transfer (µET) concept and design. (a) µET using spiral microcarriers. (b) Embryo capture and rolling motion mechanism, driven by the gliding and stepping motion of the spiral near the substrate surface. (c) Scanning electron microscope (SEM) image showing an array of spiral microcarriers. (d) MTS assay demonstrating the cell viability of Madin-Darby canine kidney (MDCK) cells co-cultured with metal-coated spirals at 37 °C for 48 h, indicating an increase in cell number on all evaluated materials and no apparent toxicity. The control group was cultured without spirals. (e) Schematic representation of the procedure’s basic principles, including capture, transport, and release of early embryos to the target region. (f) Optical images illustrating the capture, transport, and release of a dead oocyte (model cargo) to the target region in vitro. (g) Mechanical-controlled oocyte loading using a needle.

**Fig. 1b** illustrates the embryo capture mechanism and propulsion of the microrobot while moving near the substrate surface. The microrobots rotate around their short axis across a wide range of frequencies. After developing the structures, they were coated with Ti (10 nm), Fe (100 nm), and Ti (10 nm) using electron beam physical vapor deposition at a rate of 0.5–1.0 Å/s. Scanning electron microscopy (SEM) imaging was performed after coating the sample with ∼10 nm Pt to enhance conductivity and prevent charging effects (more details in the Materials and Methods) (**Fig. 1c**).

To assess the biocompatibility of the microrobots and materials used, an MTS assay (3-(4,5-dimethylthiazol-2-yl)-5-(3-carboxymethoxyphenyl)-2-(4-sulfophenyl)-2H-tetrazolium) was conducted using the Madin-Darby canine kidney (MDCK) cell line. This cell line is widely used in biomedical research to evaluate the toxicity of various compounds and their effects on cell viability and barrier integrity. The viability study was performed after co-culturing MDCK cells with metal-coated microrobots at 37°C for 48 h, showing an increase in cell numbers across all tested materials, with no signs of toxicity. A control group, cultured without microrobots, was used for comparison (**Fig. 1d**). The primary function of the spiral microrobot is to capture embryos, transport them safely using a rotating magnetic field, and release them in a controlled manner at the target location (**Fig. 1e**). For this in vitro experiment, dead mouse oocytes (model cargoes) were retrieved and carefully positioned near the microrobot opening within cell culture media. First, the microrobots were actuated to enclose the cargo, positioning it securely at their center. Then, applying a counter-rotation to the spiral microrobot enabling the controlled release of the oocyte (**Fig. 1f, movie S1**).

Another approach to load the microrobots was by using a glass capillary to guide the oocytes one by one toward the spiral’s opening and gently pushing them inside (**Fig. 1g**, **movie S1**). This method proved to be the most effective approach for loading multiple embryos into the microrobots, and we adopted this approach in next experiments (all these procedures are described in detail in the Materials and Methods).

### Propulsion in a microfluidic system cultured with endometrial cells

The experimental results of the propulsion of the spirals in different environments are presented in **Fig. 2**. The microrobots encounter several challenges in physiological conditions, particularly the actuation. These microrobots must function within the reproductive tract, which contains delicate, intricate structures, as well as viscoelastic fluids, cellular debris, and proteins. The uterus, a key organ in the reproductive system, supports and nurtures embryos, providing an optimal environment for implantation into the uterine lining. It is a complex; hollow organ filled with thick fluid and lined with a dense layer of endometrial tissue. This complexity of the uterus makes it challenging to actuate the microrobots inside this cavity and the microrobots should be able to move over endometrial epithelial cells (EECs). To better understand the motion performance of the suggested embryo carriers, and before moving to animal experiments, we performed in vitro studies, first considering confinement, then incorporating EECs cells and viscous fluids.

**Fig. 2.**
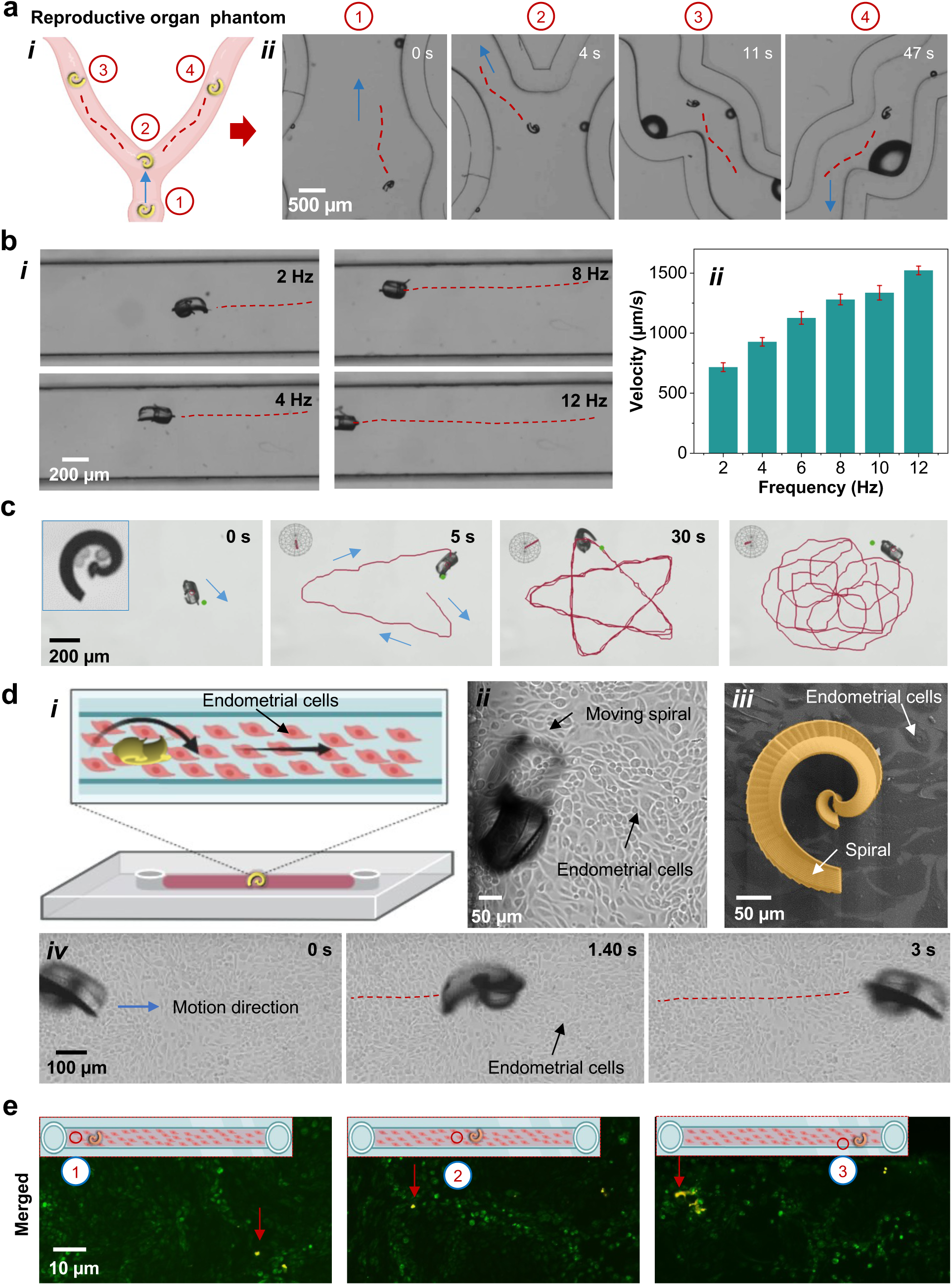
Microrobot motion performance in uterine-mimicking microfluidic systems, featuring an endometrial epithelial cell lining and ex vivo tissue. (a) A microfluidic system mimicking the female mouse reproductive tract, including the uterine body and two uterine horns (i). Spiral microrobot in an enclosed channel using an external rotating magnetic field (5 mT, 1 Hz). The single spiral was guided through the entire reproductive tract phantom, moving from positions 1 to 4 (ii). (b) Movement of the spiral in a straight PDMS channel to measure microrobot speed under a fixed field strength (5 mT) while varying frequency from 2 to 12 Hz. The spiral’s speed increases with higher frequency (i). The microrobot’s speed increases by adjusting the frequency (ii). (c) A single spiral following star-shaped and rose-shaped trajectories under closed-loop control while carrying embryos. (d) Illustration and real images of the spiral interacting with a monolayer of endometrial epithelial cells cultured in a PDMS channel: (i) schematic representation; (ii–iii) spiral moving over the cell monolayer; (iv) time-lapse images showing spiral motion across the cultured surface. (e) Fluorescence images of live-dead staining of the cell monolayer: live cells stained green (SYBR 14), dead cells stained red (PI). Inset: images of different channel sections after spiral motion over the monolayer. Three regions were selected for quantitative live-dead analysis.

To mimic typical physiological conditions of the reproductive tracts, we first designed and fabricated a microchannel with a geometry mimicking the shape of the reproductive tract of a female mouse having a uterus body and two uterine horns (**Fig. 2a, i**). The spirals were collected and immersed in DI water inside this enclosed channel and then actuated and steered by an external rotating magnetic field (5 mT, 1 Hz). A single microrobot was moved through the whole reproductive phantom as indicated from positions 1 to 4 (**Fig. 2a, ii** and **movie S2**). We also fabricated a simple confined channel and measured the speed of such microrobots with fixed field strength and varying frequency. In this experiment, we applied a field strength of 5 mT at a frequency range from 2 and 12 Hz, to steer a single spiral in a narrow channel (**Fig. 2b, i**). The microrobot’s speed increases by adjusting the frequency as shown in **Fig. 2b, ii**.

Afterward, the motion behavior of the microrobots with and without embryos was characterized using closed-loop control with optical imaging feedback and real-time magnetic actuation (**Fig. 2c**). A customized algorithm was developed to steer and control the microrobots. The bare spirals (without embryo) were immersed inside an enclosed channel and then actuated and steered by an external rotating magnetic field (**Fig. S2, movie S3**). The motion planning determines the target location to follow a star-shape as shown in time-lapse images. The interaction between the Fe layer and the magnetic field generates torque, causing a change in the microrobot’s orientation. When a rotating magnetic field is applied, the microrobot rotates and translates perpendicular to the rotational axis, moving via a rolling-like interaction with the surface^26^. The microrobot was guided using real-time imaging feedback (see details in the Materials and Methods). The difference vector to the target location was provided to the position controller, which then calculated the magnetic field vector required to induce rolling motion in that direction, as demonstrated in the time-lapse **movie S3** (2x speed) of the closed-loop control of the microrobot. The magnetic field of 8 mT and rotational frequency of 1 Hz was generated with a commercial 8-coil setup. The different trajectories were generated to check the control of such a system by modifying the algorithm. The star-shaped tracking produced an average speed of 327.9 ±31.92 µm/s and an average tracking error of 48.7 ± 10.78 µm/s. Similarly, rose-shaped was generated with an average speed of 358.4 ± 8.78 µm/s and an average tracking error of 74.1 ± 7.20 µm/s. More examples of different shapes such as figure-of-8 and house are shown in **Fig. S3 and movie S4**.

Afterward, three embryos were captured manually as explained in the previous section, and the same parameters were applied for the closed-loop control which generated similar results as shown in **Fig. 2c**. For a star-shaped trajectory, the average speed was 362.2 ±7.36 µm/s with an average tracking error of 40.1 ± 2.34 µm/s, and for a rose-shaped 378.9 ± 65.44 µm/s and 68.1 ±40.51 µm/s respectively (**Table 1**). Overall, comparing the spirals’ average speed in cell culture media (M2 and TCM-air), a slight increase can be found when carrying the embryos, while the tracking error decreases. The additional drag that arises from the captured cargo is less of a hindrance for the spirals, as they enclose the embryos and transport them close to their center of rotation by the rolling motion.

**Table 1.** Summary of spiral (with and without embryos) speed and tracking error with closed-loop control.

|  | **Spiral type** | **Velocity ($\mu\text{m/s}$)** | **Tracking error ($\mu\text{m/s}$)** |
| --- | --- | --- | --- |
| 1 | Star-shape | $327.9 \pm 31.92$ | $48.7 \pm 10.78$ |
| 2 | Star with zygotes | $362.2 \pm 7.36$ | $40.1 \pm 2.34$ |
| 3 | Rose-shape | $358.4 \pm 8.78$ | $74.1 \pm 7.20$ |
| 4 | Rose with zygote | $378.9 \pm 65.44$ | $68.1 \pm 40.51$ |

After evaluating their in vitro motion performance, the microcarriers were steered on cell-lined microfluidic devices. Endometrial Epithelial Cells (EECs), an essential component of the uterine lining (endometrium) in mice, interact with early-stage blastocysts and play a crucial role in facilitating implantation during the development process. To mimic this physiological condition, where the uterine lining is covered with EECs, a Polydimethylsiloxane (PDMS) channel was cultured with EECs to cover the whole channel surface with a monolayer of cells (**Fig. 2d, i-iii**). For this study, primary bovine epithelial endometrial cells were cultured for mimicking conditions. Ideally, the cells should be extracted from mice; however, the process proved challenging, and the yield of cells was lower compared to bovine cells, requiring more animals to obtain sufficient cells. Ex vivo mimicking samples offer an excellent opportunity to adhere to well-established ethical guidelines based on the 3Rs principles (“Replace, Reduce, Refine”), although a balance must be struck with the number of animals needed for tissue extraction. Briefly, the PDMS channels were treated with rat tail collagen I and dried overnight (more details in the Materials and Methods). Primary bovine EECs were then seeded, and the culture media was replaced every 24 h until confluence was achieved. The spiral was collected in cell culture media and released into a cell-cultured channel. The monolayer of EECs and a single spiral on the cell surface were visualized using optical microscopy. To confirm controlled motion on the cell surface, a rotating magnetic field (5 mT, 1 Hz) was applied and the spiral moved forward and backward after several attempts as shown in time-lapse images (**Fig. 2d, iv**) **and movie S5**. It is crucial to ensure that the cells remain viable, and that the spiral’s motion does not damage the cell monolayer. To assess this, the same channels were treated with Live-Dead staining. Briefly, a two-color assay was used to evaluate cell viability. Live cells were stained with SYBR 14 (green), while dead cells were stained with Propidium Iodide (PI) (red). A drop of 1 µL of SYBR 14 (diluted 1:50 from the stock solution) and 1 µL of PI solution were added to the sample, which was then incubated for 10 minutes at 37°C. The channel was imaged from three different regions of interest (ROI) (marked as 1, 2 and 3) to observe the whole channel (**Fig. 2e**) and the excitation/emission for SYBR 14 was 488/516 nm, and for PI was 535/617 nm. The channel displays a monolayer of live cells (green), with some regions showing dead cells (yellow), even after several forward and backward motions of the spiral across the cell surface.

Ex vivo fresh uterine tissue was also used to construct a fluidic channel. A fresh uterus sample was retrieved, placed in phosphate-buffered saline (PBS), and the organ was carefully opened with scissors to access the inner tissue. The spiral was then positioned on the tissue surface, as shown in the schematic (**Fig. S6, movie S6**). After the magnetic field was applied, the spiral rolled forward on the tissue surface, as shown in the time-lapse images. These results offer promising evidence that the spirals can be effectively actuated within the in vivo uterine environment for embryo transport and release experiments.

### In vitro embryo development within the spiral microrobots

This section explores the impact of spiral material on embryo development within an enclosed spiral environment, following standard procedures. For fertilization, we used IVF-generated zygotes. Oocyte collection was timed based on hormonal induction, with animals undergoing superovulation using pregnant mare serum gonadotropin (PMSG) and human chorionic gonadotropin (HCG). The oocytes were isolated the following morning for IVF and 8 h after IVF, zygotes with visible pronuclei were selected for the experiment (more details in Materials and Methods). From this group, we sorted out 18 zygotes based on the presence of pronuclei and used them for the study.

All the zygotes were divided into six groups: G1, G2, G3, G4, G5, and G6. Groups G1 and G4 each included three zygotes as controls. Groups G2 and G5 each included three zygotes captured by the spiral microrobots, while groups G3 and G6 each included three zygotes captured by the spirals and magnetically manipulated for 1 h at room temperature (RT). The qualitative results from G1 to G3 are shown in **Fig. 3a**. The data for the groups G4, G5, and G6 can be found in Supporting Information (**Fig. S5)**. To isolate the zygotes, a solution with a mixture of hyaluronidase (HA) and lyophilized powder (800 units/mg solid) was prepared with a final concentration of 1 mg/ml in M2 cell-culture media. The M2 media was freshly prepared by following the published protocol^27^. When handling the zygotes, M2 media was used for washing and sorting. During incubation, the zygotes were cultured in KSOM. We also used this KSOM medium for the 4-days development in the spiral microcarriers.

**Fig. 3.**
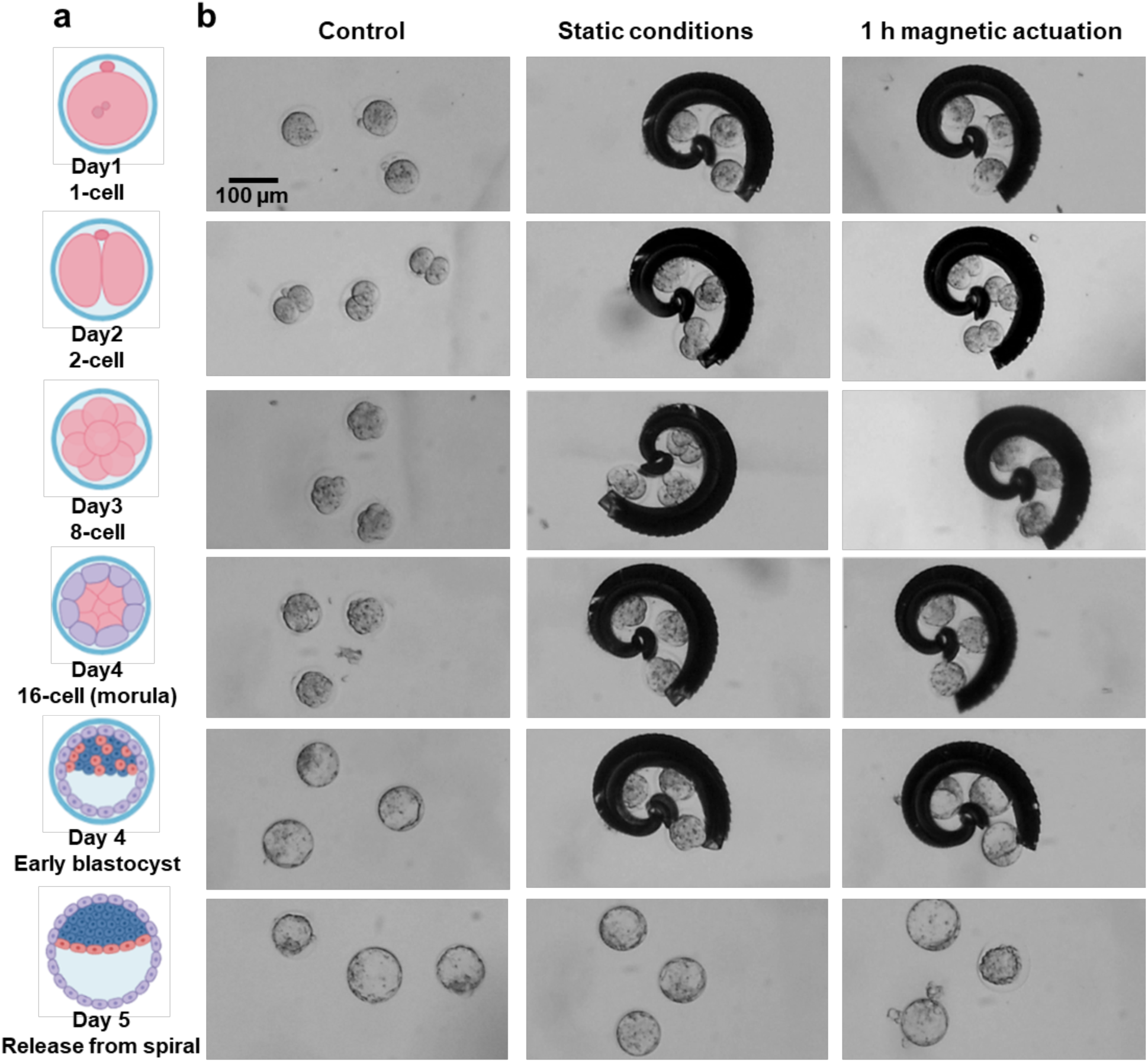
Embryo development capability – in vitro experiments. (a) Three 1-cell embryos were prepared for three different groups: G1 (control), G2 (static conditions), and G3 (magnetic actuation). The embryos were monitored until embryonic day 4 (E4) and then released from the spiral on day 5 (E5) (i-vi).

After isolation, the zygotes were maintained under physiological conditions (37°C, 5% CO_2_), while the spiral samples for groups G2 and G3 were prepared in separate Petri dishes. Three zygotes were then mechanically captured by carefully positioning them at the open end of the spiral, gently rotating the spiral for complete capture, and repeating the process for additional zygotes. Once the three zygotes were captured for group G2, the sample was imaged and returned to the incubator. A similar procedure was followed for group G3, after which the sample was magnetically treated, and motion performance was analyzed for 60– 80 minutes at RT. This procedure was performed to check whether the zygotes can develop further after going through a long handling process (**Fig. 3**). This is crucial for the real-world application of ET, where the procedure typically lasts at least 1 h, and it is essential for the embryo to remain viable for further development after implantation. A similar procedure was performed for groups G4, G5, and G6, resulting in six zygotes from each group. All samples were then optically imaged for the following four consecutive days to monitor their developmental stages. Successful progression from the 1-cell stage to early blastocyst was observed, as shown in **Fig. 3a, b** and **Fig. S5**. On day 5, early blastocysts were released from the spirals in the groups G3 and G6, and six blastocysts were carefully prepared for ET. On the transfer day, pseudo-pregnant mice were used for the unilateral surgical ET procedure into the uterine horns. A copulation plug was also detected on the morning of the transfer. After isolating the blastocysts from the spirals and washing them through 10 × 100 µl drops of M2, the embryos were transferred into the pseudo-pregnant recipient female mice. This procedure was performed under anesthesia to minimize stress and discomfort, induced by an intraperitoneal injection of a solution containing Ketamine, Xylazine, and Acepromazine. ET success and implantation were confirmed through animal dissection, showing an embryo transfer rate of 60%, which is comparable to that of conventional ET procedures in fertile mice. However, further studies are needed to evaluate subsequent embryo development (see **Fig. S6**, Supporting Information).

### Microrobots imaging and control

The imaging experiments were conducted using a dual US/PA imaging system. US provides anatomical and functional details while PA contributes to molecular information. The device was equipped with a linear array of US transducer at a central frequency of 25 MHz and fiber optic bundles on either side of the transducer for illumination. The fiber bundle was coupled to a tunable Nd: YAG laser (680 to 970 nm) with a 20 Hz repetition rate and the signals were collected by the 256-element linear array transducer. For in vitro study, a phantom setup comprised of enclosed tubing, and a water bath was prepared as shown in **Fig. 4a, i**. For the tubing phantom, a transparent intravascular polyurethane (IPU) tube (inner diameter ∼760 µm, outer diameter ∼1650 µm) was mounted in a water bath. The microrobot was inserted into the tube filled with PBS and then immersed in the phantom chamber containing DI water for better acoustic coupling. The measurements were performed with a position-fixed high-frequency transducer to avoid image distortion. Single wavelength (800 nm) imaging was performed using a 2D stepper motor and a linear translation of the transducer over the IPU tubing. The fluence was set below the Maximum Permissible Exposure (MPE) limit (20 mJ/cm^2^), followed by the safe exposure guidelines^28^. The IPU tubing features low attenuation and low front surface backscatter that provides a better US contrast making it possible to visualize a single moving spiral inside this tube phantom. The dynamic tracking of a single microrobot is shown in time-lapse images from 0 to 12 s comparing optical and US imaging (**Fig. 4a, ii, movie S7**). The yellow arrows show the position of the single-moving spiral. The spirals provided both US and PA contrast because they are coated with thin absorbing adjacent metal layers (Ti=10 nm, Fe=100 nm, Ti=10 nm). To actuate the spiral in a controlled manner, a magnetic system consisting of an integrated coil setup and an imaging system was implemented. The coils produce fields up to 50 mT in a frequency range from 0 to 200 Hz. In this experiment, we applied a field strength of 5-10 mT at a frequency range between 1-5 Hz to steer the spiral in a narrow channel. The speed of the microrobot increases by adjusting the frequency.

**Figure 4.**
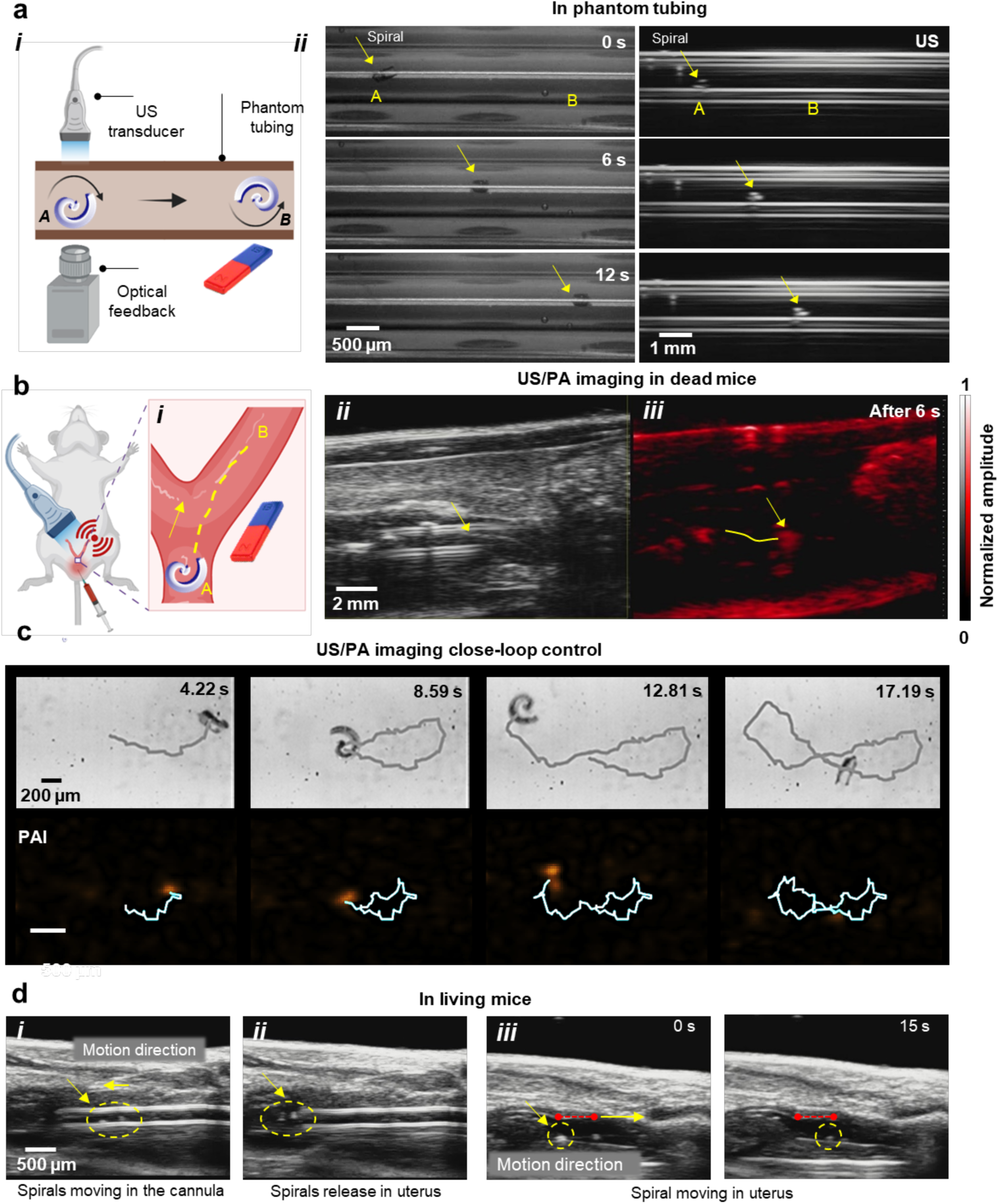
Imaging and control of spiral microrobots in different settings. (a) Schematic of dual US and optical imaging of spiral in a phantom tubing setup (i). Real-time tracking of a single moving spiral under optical and US feedback from 0 to 12 s (ii). (b) Schematic of the spiral insertion inside the uterus cavity of a dead mouse (i). The spiral is steered under a rotating magnetic field of 5 mT and 5 Hz (ii) as shown in dual US/PA imaging modes. (c) Trajectories (median-filtered over 500 ms) obtained from tracking with PA imaging and the respective optically tracked reference trajectory. (d) A cannula releasing two spiral microrobots into the mouse uterine cavity using real-time US images. After release, a single spiral is steered under a rotating magnetic field of 5-10 mT and 5 Hz as shown in time-lapse images from 0 to 15 s.

After visualizing the spiral in phantom tubing, we conducted experiments using a deceased mouse (previously used for training purposes) (**Fig. 4b, i**). The mouse was placed on a heating unit, and the catheter was inserted via the vaginal canal and cervix into the mouse uterus cavity by following safety guidelines^29^. For imaging, US gel was applied between the detector surface and the mouse’s body to match the acoustic impedance for efficient signal transfer. US mode generated a structural image, suitable to identify the tubing structure, and the PA was used to monitor the displacement of the spiral, by detecting its light absorption properties. Both images were acquired in real-time after tracking the spiral in the uterus while the spiral was inside the catheter for smooth motion performance and unhindered imaging performance as shown in time-lapse images from 0 to 6 s with clear localization of the phantom tube (**Fig. 4b, ii-iii**). The dual US/PA imaging feature is highly beneficial for deep tissue imaging, where US provides tissue background morphology, while PA imaging offers contrast based on the spiral material’s properties. The precise steering of the spiral was captured under a rotating magnetic field of 5 mT and 5 Hz, with controlled motion performance in both forward and backward directions. By leveraging the anatomical features of US, the catheter’s position inside the uterine cavity was determined, and a single spiral was steered in real-time (**movie S8**). The PA images acquired show amplitude values of the uterine cavity with the spiral microrobot, while the US images highlight tissue boundaries and morphology.

After successfully visualizing the spiral microrobot in vitro and deceased animal settings, we proceeded to investigate real-time monitoring of the moving spiral microrobot in vivo, within the uterine cavity. The in vivo experiments were performed under animal handling license number: TVV 5712021, held by the authors, ensuring the approved design of the microrobot and biocompatibility of the employed materials. This study was conducted using the same device and imaging settings as in the previous experiments. The mouse was anesthetized with an inhalant mixture of isoflurane and oxygen gas (ranging from 2.5 to 4% isoflurane), and the spirals were injected into the uterus (as detailed in the Materials and Methods). Additionally, the mouse received subcutaneous treatment with Metamizole at a dose of 0.1 ml/10 g as an analgesic, ensuring animal welfare while avoiding additional surgical stress.

Under anesthesia, the mouse uterus was catheterized by following previous safety guidelines^29^. Before starting the experiment, the physiological parameters of the mouse were monitored (**Fig. S7**). As preliminary test for closed-loop control using an imaging modality suitable for in vivo experiments, we established an experimental setup enabling simultaneous optical and PA imaging while performing closed-loop control. **Fig. 4c** illustrates an 8-shaped trajectory of a spiral microrobot captured in both optical and PA imaging modes (**movie S9**). The integration of machine learning algorithms enhances tracking efficiency and, in the future, facilitates trajectory planning and control, as reported by our group^30^.

Consequently, two spirals were collected with PBS in a catheter, which was then inserted via the vaginal canal and cervix into the uterine cavity of the anesthetized mouse, all while being monitored in vivo through US imaging (**Fig. 4d, i-ii**). Both moving spirals inside the catheter were visualized in real-time under magnetic field guidance. Upon reaching the intersection of the uterus and uterine horn, the spirals were released into the uterine body, as shown near the catheter edge. After release, the catheter was carefully retrieved, and the spirals began moving within the uterine cavity (**Fig. 4d, iii), movie S10**). Given that the spiral is ∼300 µm in size, stand-alone US imaging was sufficient to visualize a single moving spiral in vivo under a rotating magnetic field of 5-10 mT and 5 Hz. The spiral was moved within the complex uterine environment, a crucial step for vivo embryo transport. The displacement of the spiral was estimated to be ∼2.5 mm over a time interval of 15 seconds.

### Investigation of Microrobot–Tissue Interactions

To assess the immediate uterine response to µET, we used an independent cohort of mice that underwent the identical spiral-guided embryo-transfer protocol described above. Each animal was humanely euthanized immediately after the ET had been completed (0 h time-point). Reproductive tracts were washed in PBS, embedded in OCT, and frozen at - 20 °C for sectioning. Hematoxylin and Eosin (H&E) staining was performed to assess tissue morphology and evaluate the effects of the spiral microrobots on the uterine environment. Hematoxylin, which stains nucleic acids, produced a deep blue-purple color, while Eosin, which stains proteins nonspecifically, resulted in varying shades of pink in the cytoplasm and extracellular matrix. In typical tissue samples, the nuclei appeared blue, while the cytoplasm and extracellular matrix were stained pink. The reproductive organ samples were first washed in PBS and carefully fixed in cryomolds, then covered with optimum cutting temperature (OCT) compound at -20°C. Using a cryostat, we sectioned the OCT-embedded samples at 7 µm thickness for further analysis (more details are found in the Materials and Methods). Because the histology was performed immediate post-transfer, the sections represent the acute tissue response to the procedure. No significant signs of damage or abnormalities were observed in the uterine tissue. The morphology of the uterine lining appeared normal, with no indications of inflammation or adverse tissue response, suggesting that the spirals, along with their approx. 1h handling, did not adversely affect the development or integrity of the uterine tissue, compared to non-microrobot treated tissue (F**ig. S8**). These observations support the safety and biocompatibility of the microrobot material and handling conditions for potential use ET procedures (**Fig. 5a, i-iv**).

**Fig. 5.**
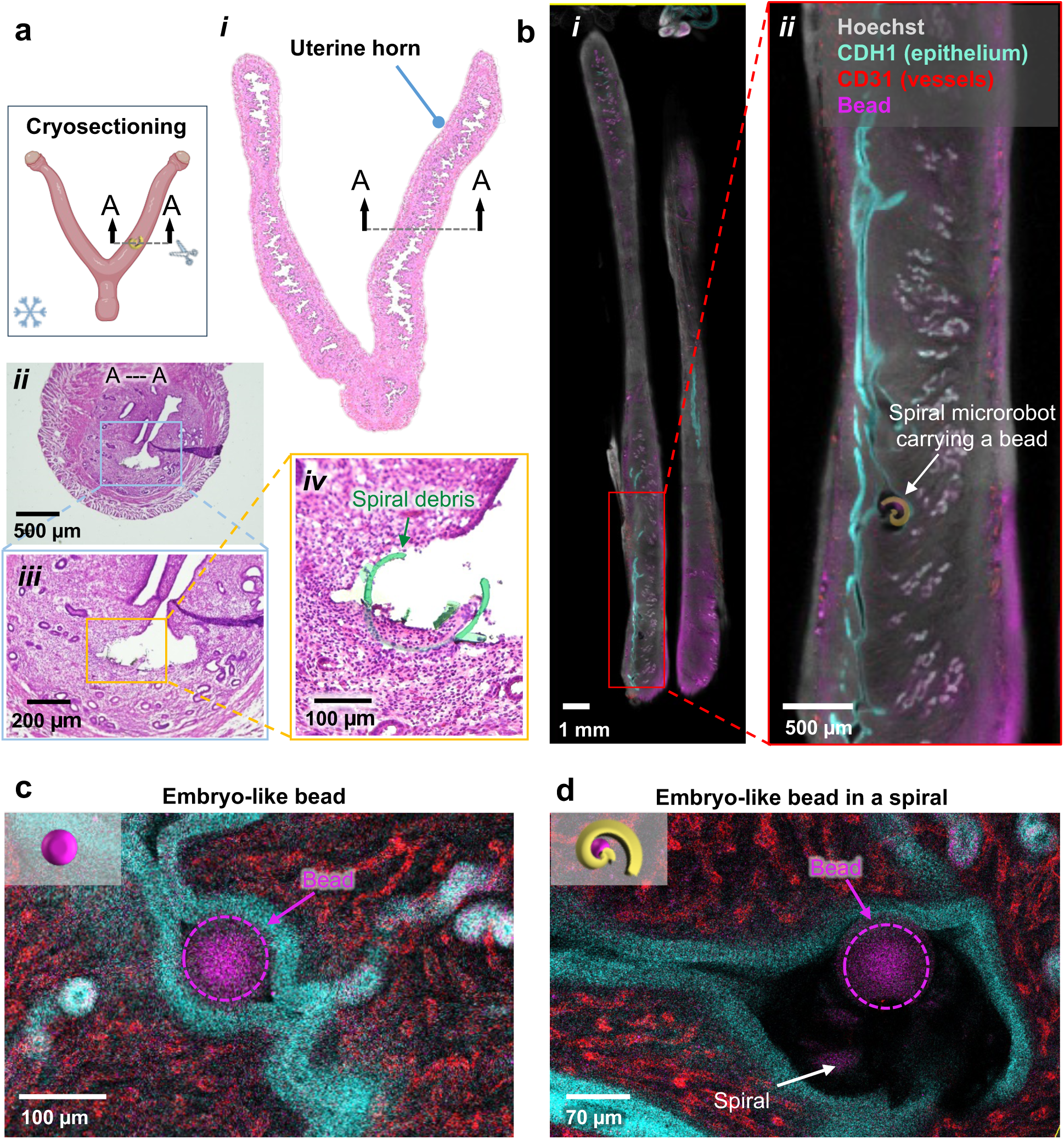
Visualization of the spiral loaded with a pseudo-embryo in the mouse uterus. (a) The whole reproductive organ harvested from the swimming microrobot-administrated mice for hematoxylin and eosin (H&E) staining analysis. (b) Confocal microscopy images of a slice view of both the uterine horns from a pseudo-pregnant mouse injected with beads and spirals loaded with beads at GD2 18:00h and evaluated at GD3 12:00h. (i) highlights the whole length uterus carrying spiral with the beads and (ii) highlights the microrobots with beads in a higher magnification image. (c) An embryo-sized bead displays a closed lumen surrounding the bead. (d) A spiral with beads showing a closed lumen surrounding the spiral. Grey: Hoechst (stains nuclei); Blue: CDH1 (stains epithelium), Pink: FOXA2 (stains glands and autofluorescence highlights beads); Red (stains blood vessels).

Finally, to assess the ability of the spiral microrobot to transport embryo-like objects in living mice, we employed whole-mount immunofluorescence and confocal imaging to analyze the microcarrier’s location within the lumen (**Fig. 5b, i-ii**). The transferred microcarrier, containing the beads, was detected in the uterus of the pregnant mice at gestational day (GD) 3, at 12:00h. The spiral microrobots were visible when the tissue was excited with a 405 nm laser, while the beads were visible under excitation with a 555 nm laser. This visibility was likely due to the non-specific binding of Hoechst (a blue DNA-binding dye) that adsorbs to exposed surfaces of the microrobots and the donkey anti-rabbit 555 secondary antibody tagged with Alexa Fluor 555, providing red fluorescence to the beads. No signal was observed when the microrobots or beads were exposed to the laser in the absence of fluorophores (data not shown). Microcarriers with beads or beads alone were found in the uterine lumen, as observed by CDH1 staining E-cadherin outlining the epithelial lumen (**Fig. 5c, d, movie S11**) surrounding the objects. At this time of mouse pregnancy, fluid in the uterine lumen is resorbed into the stroma and luminal closure occurs^31–33^. The lumen is observed to be closed around the bead or the microrobot carrying the bead. CD31 staining an endothelial marker used to verify vascular integrity reveals similar architecture of the vessels in the subepithelial stroma, suggesting that the bead or the microcarrier carrying the bead does not significantly alter the uterine architecture.

### Biodegradable Gelatin-Based Microrobots Toward Clinical Translation

While our initial demonstrations used a durable polymer (Ormocomp) to ensure robust performance, a long-term goal is to make the microrobots biodegradable so that they do not need to be retrieved after use. Moreover, by employing biosourced or naturally derived materials, we can further enhance biocompatibility and take advantage of advantageous properties—such as mild antioxidant effects and improved mechanical compliance, that may benefit embryo handling. For instance, certain natural polymers (like gelatin) can exhibit protective qualities and gentler surfaces, thereby reducing mechanical stress on delicate cells or tissues.

The photoresist polymer we employed, although biocompatible in the short term, does not naturally break down in the body, its lack of biodegradability could pose a hurdle for safe clinical application since any device remaining in the uterus after embryo delivery would ideally dissolve away harmlessly^34^. To address this, we explored the use of gelatin, a widely used biocompatible and biodegradable material, as an alternative structural material for the spirals. Gelatin is a natural polymer derived from collagen that can be cured into a hydrogel and is known to gradually degrade in the presence of bodily enzymes.

In preliminary experiments, we successfully fabricated spiral microrobots out of gelatin with a similar geometry to the original Ormocomp spirals (**Fig. 6a–c**). We incorporated magnetic microparticles (iron oxide-based) into the gelatin matrix so that the resulting gelatin spirals could still be actuated by external magnetic fields (much like the metal-coated polymer versions). The fabricated gelatin spirals were slightly more compliant (softer) than their polymer counterparts, as confirmed by compression tests (**Fig. 6d**). This increased flexibility can be advantageous, as a softer device would exert gentler forces on an embryo or on uterine tissue. We tested the magnetization and found that the embedded particles provided sufficient magnetic responsiveness, we were able to maneuver the gelatin spirals under our closed-loop magnetic control system (**Fig. 6e, movie S12**).

**Fig. 6.**
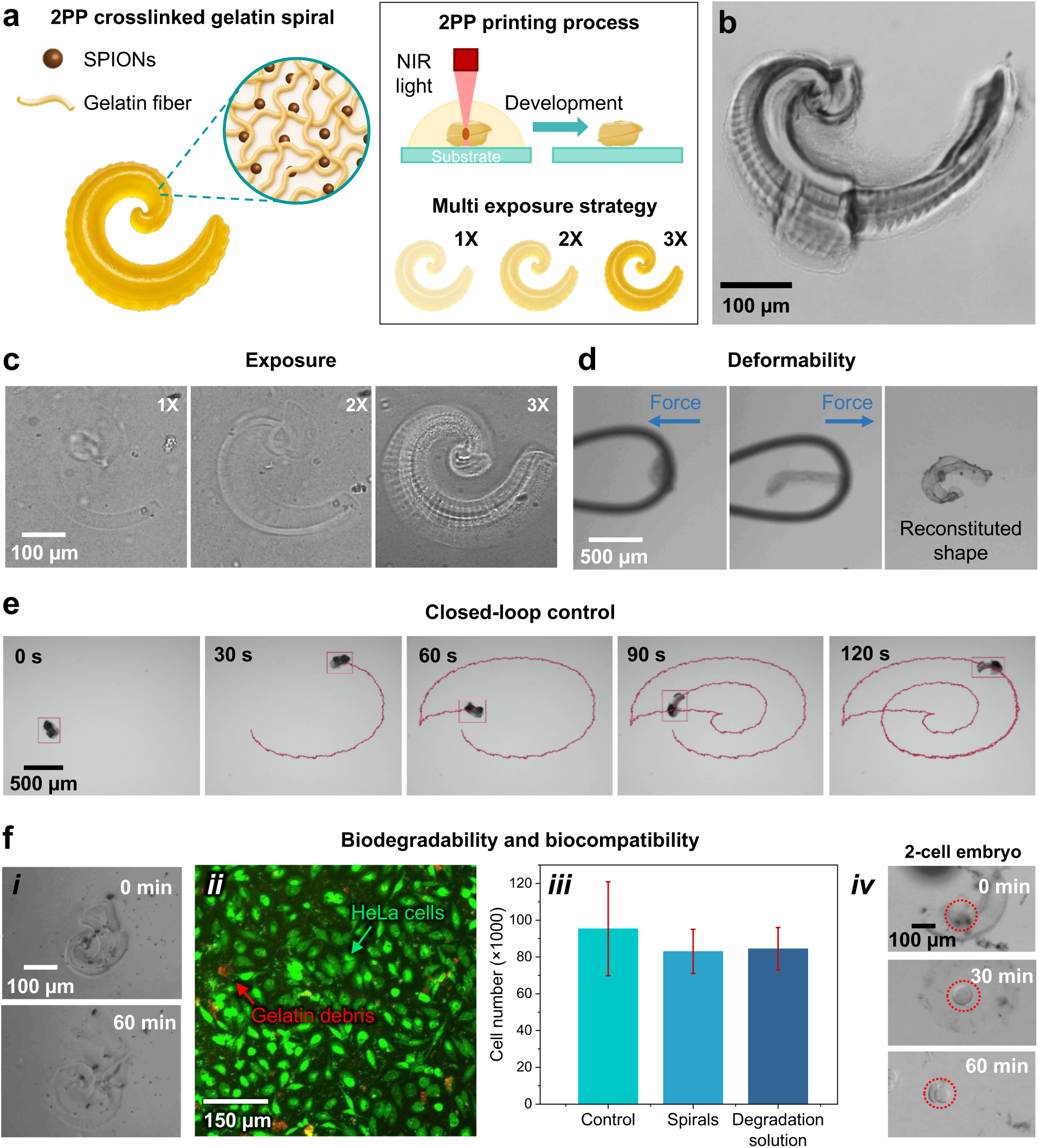
Gelatine-based spiral microrobots. (a) Schematic representation of gelatin-based microrobots embedded with SPION particles, illustrating the effects of different light exposure times. (b) The resulting gelatin-spiral microrobot. (c) Optical microscopy images of the exposed photoresist fabricated at a scanning speed of 1000 µm/sec with 60% laser power (approximately 70 mW). The labels 1X, 2X, and 3X indicate repeated exposures under the same conditions. (d) Mechanical testing of the spiral microrobot, demonstrating its high deformability and shape recovery after the applied force is removed. (e) Time-lapse images showing the complex spiral-like trajectory of the microrobot under optical imaging-based closed-loop control, demonstrating its precise maneuverability. (f) Biodegradability evaluation of spiral microrobots in the presence of gelatinase A (MMP-2) with a concentration of 5 µg/mL, over a 60-minute period (i). Live/dead staining of MDCK cells following microrobot degradation (ii), along with a quantitative comparison of cell numbers between the control group, cells cultured with spiral microrobots for 1 hour, and cells exposed to microrobot degradation products (iii). Additionally, a preliminary assessment of embryo development from the 1-cell to 2-cell stage was conducted while the microrobot degraded over a 1-hour period (iv).

Crucially, the gelatin microrobots exhibited the desired degradation behavior. In the presence of a proteolytic enzyme (we used gelatinase to simulate conditions in the uterus), the gelatin spirals gradually dissolved, breaking apart over the course of tens of minutes to a few hours depending on enzyme concentration. The degradation left behind only microscopic magnetic particles and soluble gelatin fragments, which are benign (**Fig. 6f, i**). We verified via cell culture tests that the gelatin spirals were non-toxic both before and after degradation: MDCK cells exposed to intact gelatin microrobots or to the products of completely degraded microrobots showed no reduction in viability compared to control cells (**Fig. 6f, ii-iii**). Finally, we conducted a small-scale embryo culture experiment with the gelatin microrobots. We placed mouse zygotes inside gelatin spirals and cultured them for four days, similar to our earlier procedure. The embryos developed into blastocysts on schedule, with developmental rates comparable to control embryos cultured without spirals (**Fig. 6f, iv**). This indicates that the gelatin itself (and the embedded magnetic particles) did not impair embryo development, and it suggests that a switch to a biodegradable scaffold is feasible without sacrificing functionality.

These results with gelatin are preliminary but promising. They open the door to future biodegradable microrobots that could potentially be left inside the body after delivering an embryo, whereupon they would simply break down and be resorbed, eliminating the need for any retrieval procedure.

## DISCUSSION

This study highlights the potential of magnetically actuated microrobots for enabling in vivo µET and provides key insights into several critical aspects of this technology. First, we demonstrate the efficient propulsion capabilities of spiral microrobots, showing their ability to navigate complex tissue surfaces and narrow channels, essential for replicating physiological conditions. Second, we highlight the importance of integrating a closed-loop control system for tracking microrobots in deep tissue, enabling precise and controlled navigation. Third, we demonstrate the unique ability of spiral-shaped microrobots to securely transport cellular cargo during magnetic manipulation, allowing the capture of single or multiple zygotes and supporting their subsequent development within the microrobot using only a weak external magnetic field. This capability provides a controlled environment for continuous monitoring of embryo developmental potential. Finally, we demonstrated the non-invasive delivery of embryos, previously cultured and magnetically manipulated in vitro, to the implantation site in vivo, achieving successful embryo transfer and providing initial evidence of embryo implantation.

These findings offer significant promise for advancing assisted reproductive technologies, particularly microrobot-assisted endometrial implantation and uterine transfer. Moreover, deep tissue imaging of spiral microrobots within the mouse uterus represents an important step toward in vivo assisted reproduction and the broader field of reproductive medicine.

We have also qualitatively assessed potential cytotoxicity and inflammatory effects by retrieving uterine tissues after microrobot transfer and magnetic manipulation, finding no evident adverse effects. Furthermore, our results demonstrate the feasibility of loading embryo-sized beads onto microcarriers and successfully transferring them into the mouse uterine tube during the pre-implantation phase of pregnancy. The epithelial and blood vessel architecture appeared intact, suggesting that the beads do not disrupt the structure of the uterus.

These initial studies lay the groundwork for further exploration of magnetically driven microrobots in the transport, delivery, and implantation of embryos. Future statistical studies in living mice will evaluate whether these microrobots can achieve higher embryo implantation rates compared to conventional ET. Upcoming investigations will focus not only on improving the precision of embryo localization and delivery but also on using microrobots as mechanical anchors to assist implantation and enhance endometrial receptivity by mitigating potential immune reactions that could reject the embryo. Additionally, microrobots will serve as exploratory tools for developing physical and biochemical strategies to further improve embryo transfer, monitoring, and implantation.

To translate this technology to human applications, several technical limitations need to be addressed, despite the ethical discussions surrounding the use of medical microrobots in the coming years. One key challenge is imaging penetration depth, which currently reaches approximately 2-3 cm with micrometric resolution. However, extending this depth further may compromise both spatial and temporal resolution. Other techniques, such as X-ray fluoroscopy or MRI, could enable deeper real-time imaging; however, more comprehensive studies are needed to determine whether X-ray exposure or stronger magnetic fields affect gamete or embryo function and development.

An essential step toward clinical µET is eliminating the need to retrieve the carrier after embryo release. Non-degradable photoresists, though excellent for early proof-of-concept work, are sub-optimal for long-term safety. Our preliminary switch to a biosourced material, gelatin, demonstrates that spiral microrobots can be manufactured from a natural, enzymatically resorbable material without compromising magnetic control and enhancing embryo compatibility. Because gelatin hydrogels are softer than conventional resists, they are likely to impart lower mechanical stress on both the embryo and uterine epithelium, and their intrinsic antioxidant and cell-adhesive motifs may further promote a receptive micro-environment. These findings open a path for the development of fully bioresorbable µET carriers; future studies should quantify in vivo degradation kinetics, examine possible immune modulation, and test embryo-to-birth outcomes after gelatin-spiral µET. An alternative approach would be the use of magnetically tethered microrobots with a hollow channel or microgrippers, allowing the release of embryos or therapeutic cargo while enabling retrieval from the outside.

In conclusion, this study introduces a novel methodology leveraging microrobotics for the non-invasive in vivo transfer of embryos to the implantation site. The microrobot design, in combination with the integrated imaging setup, presents a promising approach for enhancing supervised in vivo assisted reproduction techniques. However, additional statistical analysis on embryo implantation rates using microrobots is necessary, with the ultimate goal of demonstrating the successful transport of early embryos to the fallopian tubes, in cases of recurrent embryo implantation failure. Furthermore, it is crucial to assess the health and fertility outcomes of the offspring following microrobotic-assisted treatment we aim to further develop and evaluate the biodegradable gelatin-based microrobots, alongside exploring implantation rates in obesity and aging mouse models, as well as in mice with endometrial receptivity deficiencies, which are relevant for infertility research. Moreover, this technology shows great potential for expanding into applications for treating urinary tract diseases, bladder cancer, and infections, offering a promising direction for future medical innovations.

## MATERIALS AND METHODS

### Fabrication of microrobots

Spiral-shaped microrobots were designed and programmed in a general writing language editor (DeScribe, Nanoscribe GmbH, Germany) for direct laser writing (DLW) as shown previously^16^. A 3D laser lithography technology based on two-photon absorption and polymerization of negative tone photoresist (Photonic Professional GT 3D, Nanoscribe GmbH, Germany). Ormocomp (Micro Resist Technology GmbH, Germany) was employed as a photoresist, drop-cast onto fused silica substrates, and developed for 10 min (mr-Dev 600, Micro Resist Technology GmbH, Germany) after DLW-patterning according to the programmed scripts. The developed samples were immersed in isopropanol and dried in a critical point drying machine (EM CPD300, Leica Microsystems GmbH, Germany). Then, the helices were coated with 10 nm Ti, 100 nm Fe, and 10 nm Ti by electron beam evaporation (PLASSYS Bestek Ltd., France).

### MTT assay to assess microrobots biocompatibility

The MTS assay is a quantitative colorimetric method used for the rapid, sensitive, and accurate measurement of viable cells in proliferation assays (MTS Assay Kit, Abcam, ab197010). In this study, MDCK cells (∼5000 cells per well) were cultured in a 96-well microtiter plate with a final volume of 200 µL per well and incubated for 48 h at 38°C in the dark. Following incubation, 20 µL of MTS reagent was added to each well and further incubated for 2 h under standard culture conditions at 38°C. Before measuring absorbance, the plate was gently shaken, and optical density (OD) was recorded at 490 nm.

### Fabrication of microfluidic channels

Microfluidic channels were fabricated using PDMS on a glass surface and parafilm (Parafilm, Merck KGaA, Germany) sandwiched between two glass slides. Briefly, base and curing agents for the PDMS were poured into a custom-made poly(methyl methacrylate) (PMMA) mold and cured overnight. The channels obtained were then cut to fit onto glass coverslips and pierced with a 2 mm punch to obtain inlets and outlets before fixing them on the glass. Tight fixation was achieved by subjecting both surfaces, channel and glass, to O_2_ plasma (Femto system, Diener Electronic GmbH & Co. KG, Germany) for 30 s before pressing them together manually and curing the bonding for 30 min at 65 °C. Parafilm channels were fabricated by folding a strip of parafilm into three layers and cutting the desired channels into it with an electronic cutting machine (Silhouette CAMEO, Silhouette America Inc., USA). The stacks of layers with the channel shapes were then pressed and fixed between a glass slide and glass coverslip by partial melting and bonding for ∼20 s at 120 °C on a hot plate. All channels were filled with Pluronic (Pluronic F-127, Merck KGaA, Germany) solution (10 µg mL^−1^ in deionized water) and incubated overnight at 37 °C, then rinsed with deionized water and sterilized under UV light before use.

### Fabrication of microfluidic channels coated with endometrial epithelial cells

Bovine uteri were obtained from a slaughterhouse (Vorwerk Podemus e.K., Dresden, Germany), and bovine endometrial epithelial cells were isolated and cultured following the protocol described by Murillo and Muñoz^36^. Briefly, the uteri were transported in ice to the laboratory within 90 minutes after collection. The external surface was washed with 70% ethanol, and the ipsilateral horn was placed above a cooling block inside the safety bench for manipulation. The horn was opened to expose the endometrial tissue and wash it with PBS with antibiotics, then the endometrial tissue was carefully extracted. The pieces of endometrial tissue were digested for 1 hour at 37°C in a water bath. After stopping the reaction, the solution was passed through a 40 μm pore-size strainer, in which the endometrial clusters will be captured. Flip the strainer into a new tube and wash with neutralizing solution. Centrifuge the tube, discard the supernatant and resuspend the pellet into a new neutralizing solution. Centrifuge one more time, discard supernatant and resuspend in culture media. Cells were seeded in a T75 culture flask dish for 18 h; the stromal cells should be attached to the flask, and the media containing endometrial epithelial cells was removed and placed in a new flask. After each passage, the flask containing endometrial cells was checked for purity using Vimentin and Cytokeratin. Once purity was obtained, cells were used for the experiments. A 3-centimeter straight microfluidic PDMS channel was utilized for endometrial cell culture. Initially, the channels were coated with rat tail collagen type I (Serva) and allowed to dry overnight. Subsequently, the cells were seeded, and media changes were performed every 24 h until a monolayer of cells was achieved for the subsequent experiments.

### In vitro fertilization (IVF)

Mice oocytes were isolated, cultured, matured, and fertilized by IVF with mice sperm. Murine sperm and oocytes were collected from unused sources from running projects of generating mutant and rederiving mouse lines by IVF with frozen sperm in the facility. For isolation, flushing, and washing, we used M2 media; for the IVF, we used HTF media; and for incubation, we used KSOM media. These media are standards in mouse embryology. During IVF and incubation, we covered the media drops with paraffin oil. The Transgenic Core Facility at MPI-CBG Dresden holds active permissions for the work with mouse embryos and works under the principles of the 3Rs with animals living under specific pathogen-free (SPF) conditions. Before and after the microrobot experiments, the fertilized oocytes were incubated at 37 °C and 5% CO_2_ in the respective cell culture medium. The composition of M2 and KSOM media is included in the Supporting Information section of the Supplementary Materials.

### Magnetic actuation of the spiral microrobots

The rotating magnetic field for the actuation of spiral-shaped microrobots was generated using a commercial electromagnetic coil setup (MFG-100-i, Magnebotix AG, Switzerland), which was mounted onto an inverted microscope (Eclipse Ti2, Nikon Corp., Japan) to actuate and control the microrobots inside the previously described microfluidic channels under live observation and recording at 4 and 10× magnification and 10 fps (DS-Qi2 camera, Nikon Corp. Japan). The microrobots were separated from their fused silica substrate after fabrication by gentle swiping with a 10 µL pipette tip after a drop of liquid medium was placed on the respective array of microrobots on the substrate. The following liquid media were used for different experiments: murine oocyte/zygote cell culture medium (M2). The M2 was prepared according to established protocols (see IVF and Cell Culture Section). The microrobot samples were subjected to O2 plasma (Femto system, Diener Electronic GmbH & Co. KG, Germany) for 30 s to improve wetting before being suspended in the medium. The separated and suspended microrobots were transferred to the parafilm or PDMS channels and 10 µL pipette tips serving as the tubular microchannels by pipetting. Murine oocytes/zygotes were also added by pipetting with the M2 medium. The microrobots were actuated with magnetic flux densities of 1–20 mT and field rotation frequencies of 0.5–70 Hz and steered by tilting the axis of rotation of the magnetic field with a graphical user interface (Daedalus, Magnebotix AG, Switzerland) and 3D mouse (SpaceMouse, 3Dconnexion GmbH, Germany) connected to the setup.

### Closed-loop control

Having the microrobots follow a predefined trajectory in an automatic way under in vivo conditions requires closed-loop position control. We implement real-time guidance based on optical and photoacoustic imaging feedback. For the former a digital camera (a2A2590-60ucPRO, Basler AG, Germany) captures monochromatic bright-field image frames (2592 x 1944 pixels, 8 bit per pixel) at 20 Hz through an inverted microscope (Eclipse Ti2-E, Nikon Corp., Japan). Real-time photoacoustical guidance was achieved combining an open ultrasound research platform (us4R, us4us, Warsaw, Poland) for ultrasound acquisition with raw-data streaming from 256 elements, 17 MHz center frequency CMUT array (L22-8 Kolo Medical) with synchronized laser excitation (Q-TUNE-E10-SH, Quantum Light Instruments Ltd., Lithuania) at 10 Hz and a wavelength of 1053 nm. The image processing for binarization with a configurable threshold and calculation of the centroid was implemented based on CuPy ^39^ and performed on a GPU (GeForce RTX 3090, Nvidia, USA). The localized microrobots are linked between consecutive frames using TrackPy ^40^ running on CPUs (2x Intel Xeon Gold 5217, Intel, USA). The difference vector to the target location was fed to the position controller, which calculates the magnetic field vector that is generated with a commercial 8-coil setup (MFG-100-i, Magnebotix AG, Switzerland).

### US/PA imaging

The Vevo F2 LAZR-X Multi-Modal Imaging System (FUJIFILM VisualSonics, The Netherlands), which allows for dual US/PA imaging, was used for the in vitro experiments. The measurements were conducted using a 256-element linear array US transducer with a central frequency of 21 MHz. The collected signals were processed and reconstructed via onboard automated postprocessing. The system also featured fiber optic bundles positioned on either side of the US transducer for illumination. These fiber bundles were coupled to a tunable Nd:YAG laser (680–970 nm) with a 20 Hz repetition rate. For single-pulse excitation, PA images were captured at an excitation wavelength of 915 nm, with in-plane axial resolution of 75 μm and temporal resolution ranging from 5 to 20 fps.

For the experiments, commercially available methacrylate support (Vevo Phantom, FUJIFILM VisualSonics, The Netherlands) was used to mount transparent intravascular polyurethane tubing (inner diameter ≈ 380 μm, outer diameter ≈ 840 μm, SAI Infusion Technologies, USA), which served as a fluidic channel for the in vitro experiments.

Experiments in mice involved several preparatory and monitoring steps. Then, assess the health status and administer analgesia with subcutaneous Metamizol (4 ml/kg). Anesthesia was then induced with 4% isoflurane and 0.8-1 L/min O_2_ in an inhalation box, followed by maintenance with 1.5 to 2.5% isoflurane and 0.8 L/min O_2_. The depth of anesthesia was monitored through respiration, muscle relaxation, whisker movement, eyelid reflexes, and flexor reflexes. Eye ointment (Bepanthen) and a 200 µl 5% glucose solution are applied subcutaneously. The mouse was connected to sensors for continuous monitoring of heart rate (530 bpm), respiratory rate (140-150 bpm), body temperature, and mucosal color. After dry shaving the abdominal fur, US gel was applied, and a catheter was inserted into the vagina and uterus under US supervision. Vital parameters were monitored every 10-15 minutes, and the procedure did not exceed 1 hour. Then the US/PA probe was positioned on the abdominal side of the mice, while lying down on a heating plate to maintain the corporal temperature to 38 degrees. The heating plate was placed in the center of a set of electromagnetic coils (Octomag system, Magnebotix®) for the remote actuation of microrobots in living mice. The in vivo experiments were performed under animal handling license number: TVV 5712021 (Saxony, Germany).

During imaging, the US probe was manually focused on the phantom region of interest and secured in a holder. US images and movies were acquired in bright mode using the 21 MHz probe at a gain of 8 dB. Simultaneously, PA images and movies were captured using the same probe with a PA gain of 40 dB and a laser operating at a wavelength range of 785– 910 nm.

### Live dead staining

A two-color assay was used to assess cell viability. The live cells were stained with SYBR 14 (in green) and the dead cells with Propidium Iodide or PI (in red) (Invitrogen™ Catalog number L7011). We followed the protocol the company recommends; we added 1 µL of SYBR 14 1:50 dilution of the stock solution and 1 µL of the PI solution to the sample. Incubate the sample for 5-10 minutes at 37°C. The excitation/emission for SYBR 14 is 488/516 nm and PI 535/617 nm.

PDMS channels are coated with primary bovine epithelial endometrial cells. The PDMS channels were treated with Rat tail collagen I (Serva, 47254.01, Lot 191080) and let dry overnight. Primary bovine epithelial endometrial cells were seeded, and the media culture was changed every 24 h until confluency was reached.

### Histology analysis

Following vivo imaging, reproductive organs from the mice were carefully dissected for histological analysis. Hematoxylin and Eosin (H&E) staining was performed. Hematoxylin exhibits a deep blue-purple color and stains nucleic acids, while Eosin is pink and nonspecifically stains proteins. Consequently, in typically stained tissue, nuclei appear blue, whereas the cytoplasm and extracellular matrix display varying shades of pink staining. The procedure involved the following steps: Samples were washed in PBS, carefully fixed in cryomolds, then covered with optimum cutting temperature (OCT) compound (Sakura Finetek USA Inc), and subsequently frozen at -20°C. Subsequently, the OCT-embedded samples were sectioned using a “Cryostar NX70” cryostat (Thermo Fisher Scientific/ Epredia). Sections were cut to a thickness of 7 µm at a sample temperature of -17°C to - 18°C (knife temperature: -30°C) and collected onto Superfrost Plus slides. The sections were stored at -20°C. Following sectioning, the slides were prepared for staining by allowing them to air-dry for at least 1 hour at RT. Staining was conducted in cuvettes previously filled with Hematoxylin (Richard Allen Scientific, Hematoxylin 7211), Eosin (Epredia, Eosin-Y alcoholic, 71211), other required solvents, and rinse solutions. The staining sequence involved immersing slides in Hematoxylin for 5 minutes, followed by a 5-minute rinse in flowing [?] deionized (DI) water. Next, slides were immersed in Eosin for 10 seconds and rinsed again in flowing [?] DI water for 2 minutes. Dehydration was subsequently performed through sequential 2-minute immersions in 40% Ethanol, 70% Ethanol, 96% Ethanol (three times), and 100% Ethanol (twice). Slides were then cleared by immersion in Xylene for 2 minutes (twice). Finally, the stained sections were mounted using Cytoseal XYL (Thermo Scientific) and covered with coverslip.

### In vivo transfer of microcarriers with loaded beads

CD1 mice were obtained from Charles River Laboratories. For pregnancy studies, adult females aged 6 to 8 weeks were mated with fertile males, and the appearance of a vaginal plug was identified as gestational day (GD) 0.5. To visualize the transferred microrobots, the CD1 females were dissected at GD3 1200 h. Mouse studies involving the transfer of microrobots with beads in pregnant mice were approved by the Michigan State University Institutional Animal Care and Use Committee. To prepare Concanavalin A (Con A) beads (1003090735, Sigma), we washed 20uL of the beads with saline 4x times 24 h before transferring them into the microrobots. The beads were allowed to settle at the bottom of the microfuge tube each time they were washed. Finally, the beads were stored in saline at 4_°_C. A drop of saline with about five beads and the microrobot was added to the petri dish. The microrobot’s movement was controlled using a magnet. One or two beads were loaded into a glass mouth pipette. After positioning the microrobot, the beads were loaded into it, similar to the method described by Schwarz et al. (2020). After loading the beads into the microrobots, the microrobot was picked up using the small cannula of the non-surgical ET (NSET) device (60010, ParaTechs) and transferred into pregnant mice at GD2 1800h. At GD3 1200h, the mice were dissected for analysis.

### Whole-mount immunofluorescence

The whole-mount immunofluorescence staining was performed on whole tissue as described previously (Arora et al., 2016; Flores et al., 2020; Madhavan et al., 2022). The dissected uterus was fixed in DMSO: methanol (1:4). The uterus was then hydrated in methanol:PBT (1% Triton X-100 in PBS) (1:1) for 15 minutes, followed by a 15-minute wash in PBT. Next, the uterus was incubated in a blocking solution (2% powdered milk in PBT) for 2 h at RT, followed by seven nights of incubation at 4°C with primary antibodies diluted in the blocking solution. After incubation, the uterus was washed with 1% PBT for 2X 15 minutes and 4X 45 minutes, followed by incubation with secondary antibodies at 4°C for three nights. Following this, the uterus was washed with 1% PBT for 1X 15 minutes and 3X 45 minutes before being incubated at 4°C overnight with 3% H_2_O_2_ diluted in methanol. Finally, the uteri were washed with 100% methanol for 2X 15 minutes and 1X 60 minutes, followed by tissue clearing with benzyl alcohol: benzyl benzoate (1:2) (Sigma-Aldrich, 108006, B6630) overnight. Primary antibodies include rat anti-CDH1 (M108, Takara Biosciences; 1:500), rabbit anti-FOXA2 (ab108422, Abcam; 1:500), and Armenian-hamster anti-CD31 (AB_2161039, DSHB; 1:200). Alexa Flour-conjugated secondary antibodies included donkey anti-rat 488 (A21208, Invitrogen; 1:500), donkey anti-rabbit 555 (A31572, Invitrogen; 1:500), goat anti-Armenian hamster 647 (A78967, Fisher Scientific; 1:500), and Hoechst (Sigma Aldrich, B2261).

### Confocal microscopy

After performing whole mount staining, the uterus was imaged using a Leica TCS SP8 X Confocal Laser Scanning Microscope System with a white-light laser. Utilizing a 10x air objective, the entire uterine tissue was imaged with a 7.0 um Z stack (Madhavan et al., 2022). Images were merged with Leica software LASX version 3.5.5 and saved as .LIF files. The microrobots were visible when the tissue was excited by the 405nm laser, and the beads were visible when the tissue was excited with 555nm laser. Image analysis was conducted using Imaris software v9.2.1 (Bitplane). The confocal image file (.LIF) was imported into the Surpass mode of Imaris. microrobots (405nm laser) and ConA beads (555nm laser) were visible, and 3D surfaces were reconstructed.

### Statistics

Movies and images of the microrobot experiments were analyzed with Fiji software and microrobot velocities were measured with the MTrackJ plugin by Erik Meijering (https://imagescience.org/ meijering/software/mtrackj/). Each helix or spiral-shaped microrobot was analyzed at different actuation frequencies, providing multiple tracks with multiple track points, and therefore average velocities with standard deviations for multiple frequencies. The respective maximum velocities of several spirals (n = 9), obtained at different frequencies, are averaged (with standard deviation) and summarized in Figure 3A. Cases of spirals (n = 4) where the cargo was transported by the same microrobot and distinctive tracks before and after cargo coupling were successfully recorded are summarized analogously in Figure 4C, normalized by setting all individual velocities before coupling to one. The standard deviations displayed in both graphs permit the obtained calculations.

## Acknowledgments

The authors thank Dr. Lukas Schwarz for spiral design and Ms. Cornelia Geringswald for 3D printing. We thank the slaughterhouse Podemus workers and Ms. F. Hebenstreit for providing bovine female reproductive organs to extract endometrial cells. We should also thank Katrin Reppe, from MPI-CBG (Dresden, Germany) for the support on the animal license preparation.

## Funding

1. M. Medina-Sánchez thanks the financial support received from the European Union’s Horizon 2020 research and innovation program (ERC Starting Grant Nr. 853609). And the HORIZON-MSCA-2022-COFUND-101126600-SmartBRAIN3. This work was supported in part by NIH R01HD109152 to RA and T32HD087166 (to H.R.K). And this work was also supported by the Spanish Ministry of Science, Innovation, and Universities through the ‘Generación de Conocimiento’ program (Project reference: MICIU/AEI/ 10.13039/501100011033)

## Author contributions

Conceptualization: M.M-S. Methodology: A.A., D.C.R., Z.C., and M.M.S. Software: Ri.N. Validation: A.A., D.C.R., Z.C, and M.M-S. Formal analysis: A.A, D.C.R, Z.C, and M.M-S. Investigation: A.A., D.C.R., Z.C., C.R., Ro.N., Ri.N., H.R.K, R.A. and M.M-S. Resources: Ro.N., R.A., and M.M-S. Data curation: A.A., D.C.R., Z.C., and M.M-S. Writing—original draft: A.A and M.M-S. Writing—review and editing: All authors. Visualization: A.A., D.C.R., Z.C, and M.M-S. Supervision: M.M-S. Project administration: M.M-S. Funding acquisition: M.M-S.

## Competing interests

The authors declare no competing interests.

## Data and materials availability

All data are provided in the manuscript and the Supplementary Materials.

## Notes

### Competing Interest Statement

The authors have declared no competing interest.

