## Supporting Information for "Magnetically Controlled Microrobots for In Vivo Non-Invasive Embryo Transfer"

[1]NanoBiosystems Group, CIC nanoGUNE, San Sebastián, Spain.
[2]Institute for Solid State and Materials Research (IFW), Dresden, Germany.
[3]Chair of Micro- and NanoSystems, Center for Molecular Bioengineering (B CUBE), Dresden University of Technology, Dresden, Germany.
[4]Transgenic Core Facility, Max Planck Institute of Molecular Cell Biology and Genetics (MPI-CBG), Dresden, Germany
[5]Department of Obstetrics Gynecology and Reproductive Biology, Michigan State University, United States.
[6]Vodafone Chair Mobile Communications Systems, Department of Electrical Engineering, Technische Universität Dresden (current affiliation)
[7]IKERBASQUE, Basque Foundation for Science, Bilbao, Spain.

[ζ]Shared first authorship

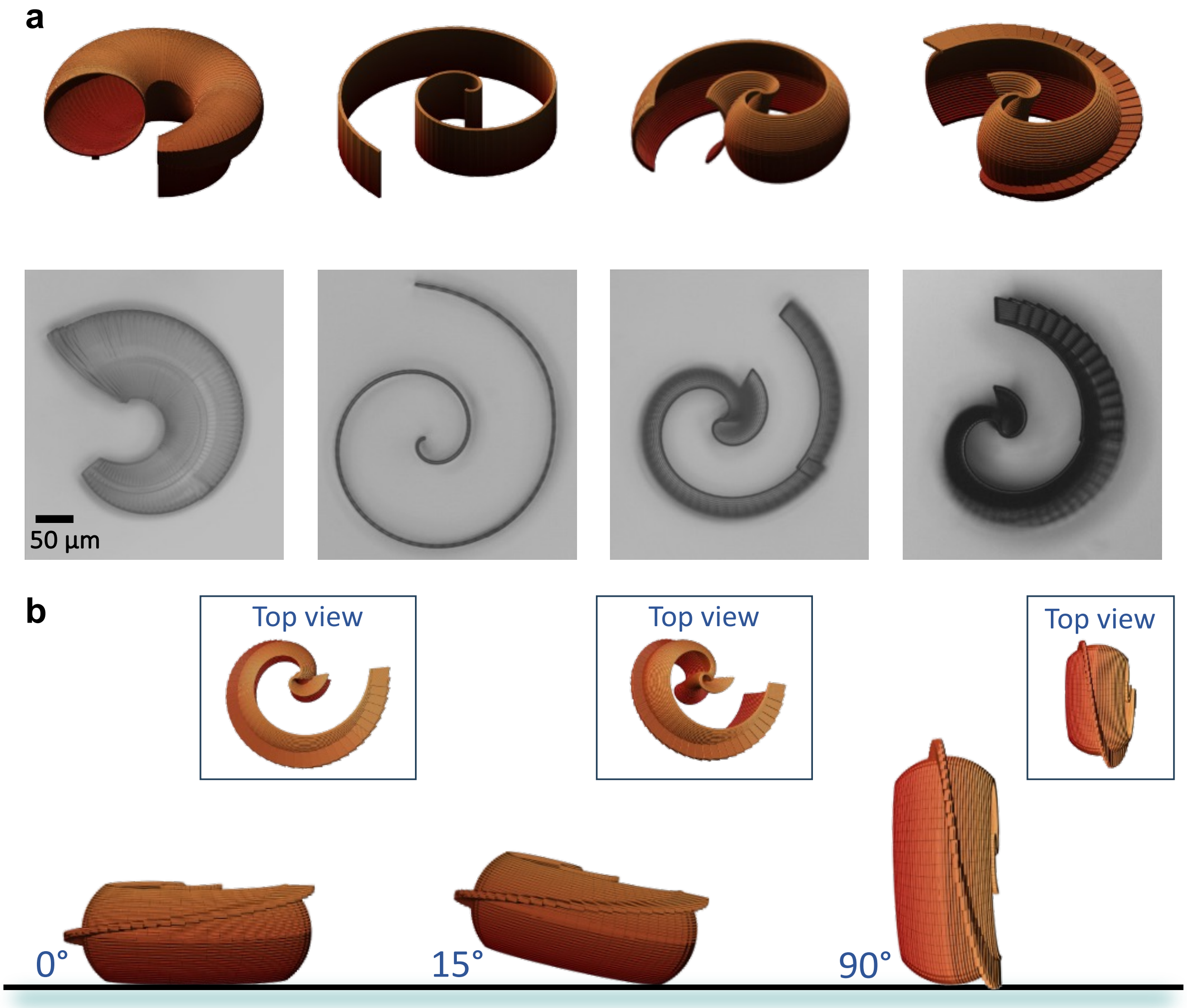


**Fig. S1. Spiral microrobot design opmitzation:** (a) Iterative spiral designs evaluated for enhanced performance optimization, and (b) locomotion principle of the optimized spiral microrobot

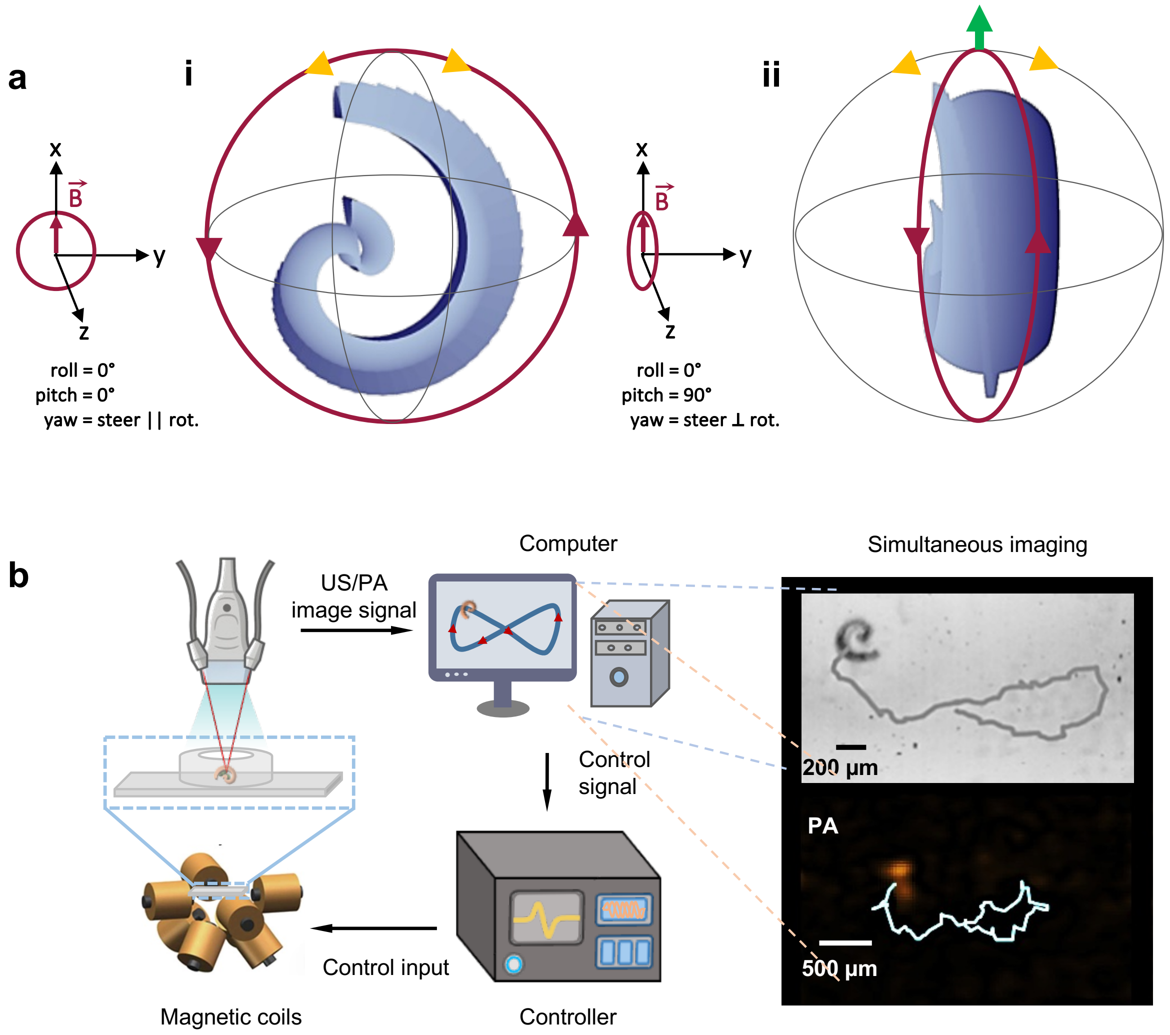


**Fig. S2. Locomotion principle and actuation platform:** (a) Rolling mechanisms of spiral microrobots, (b)Setup for conducting an in-chip parametric study of spiral microrobot locomotion under varying magnetic fields.

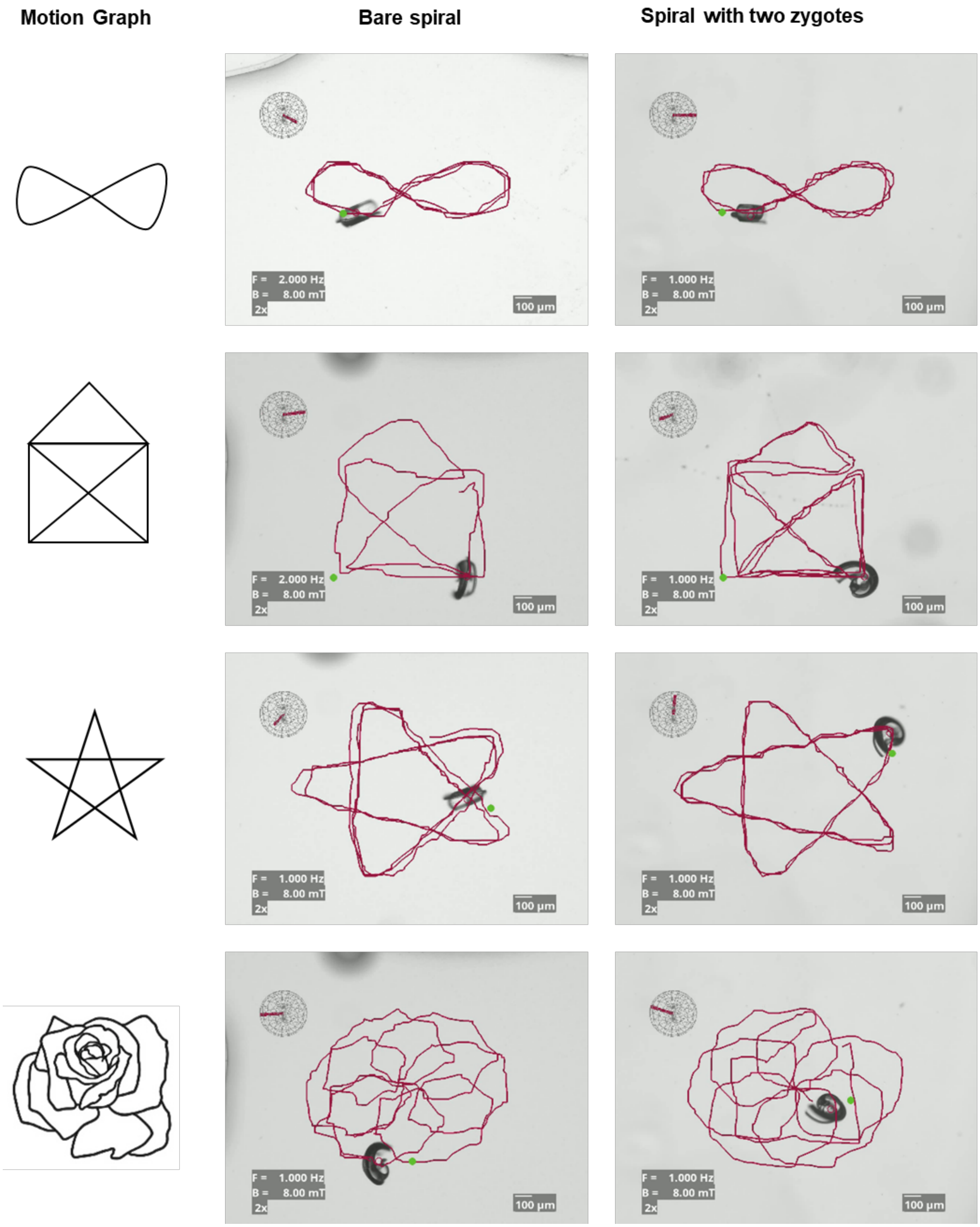


**Fig. S3.** Closed-loop control of spiral microrobots, with and without zygotes, following various predefined trajectories.

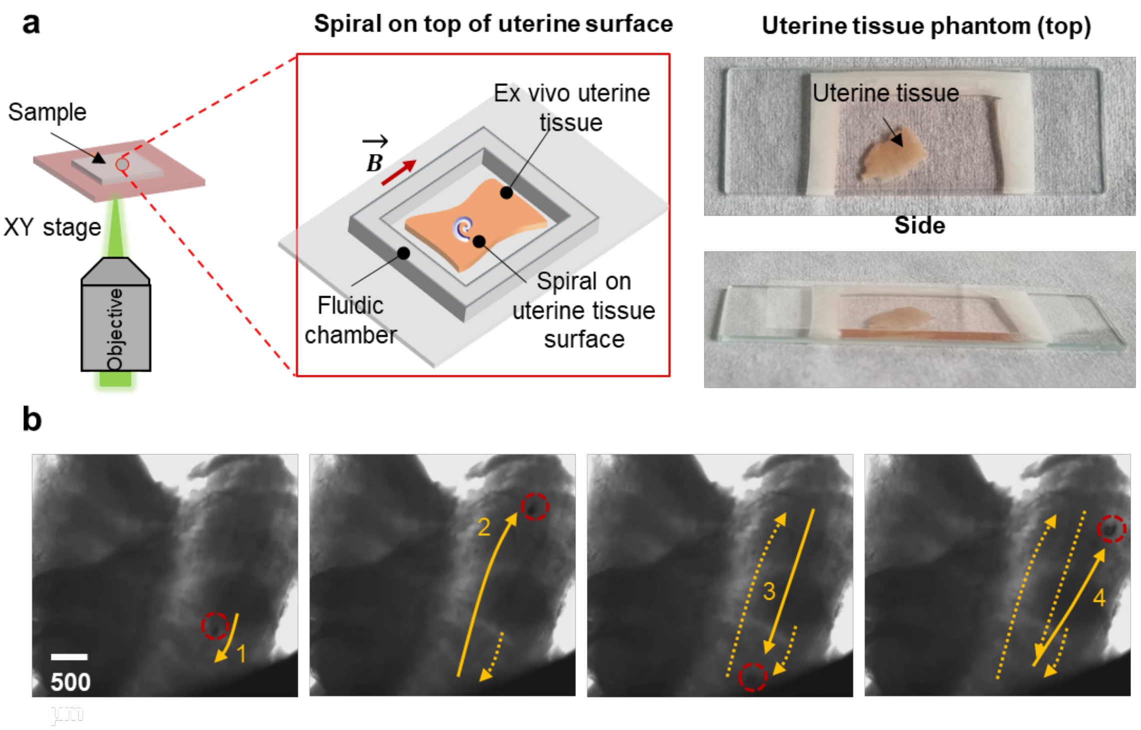


**Fig. S4.** Spiral microrobot moving on ex vivo tissue. (a) Experimental setup for optical tracking and control of the microrobot using a parafilm chip, with the bottom layer covered by flat ex vivo uterine tissue. (b) Trajectory of the spiral microrobot over time captured by optical microscopy.

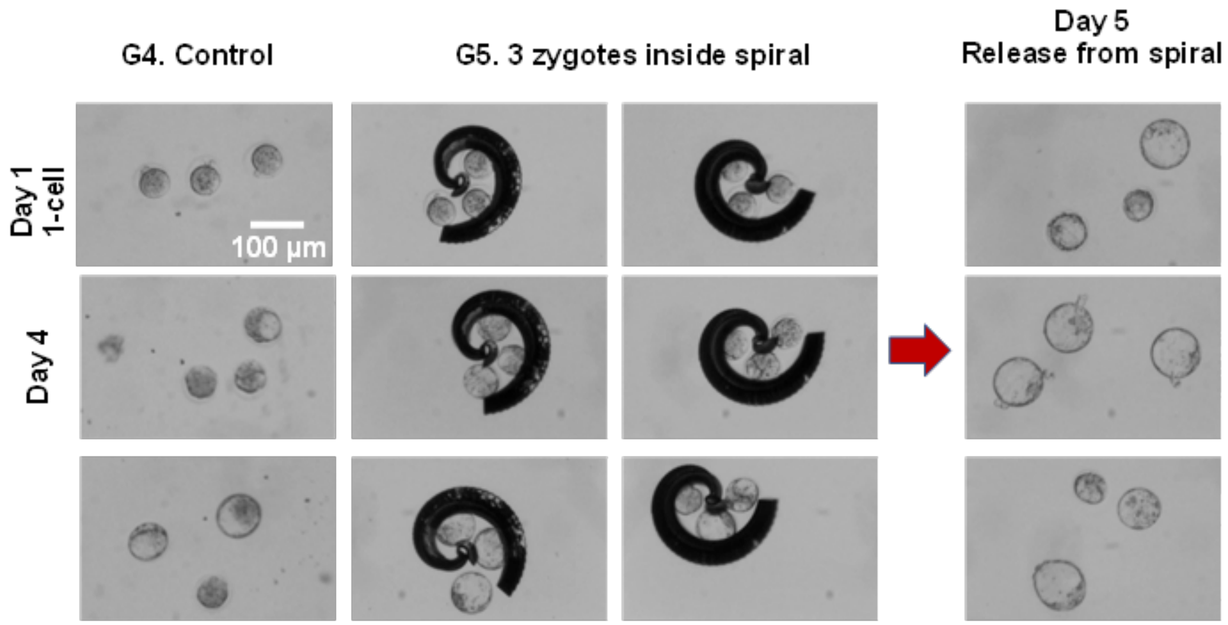


**Fig. S5.** Tracking embryo development from the 2-cell stage to blastocyst, followed by their release for subsequent embryo transfer and implantation studies.

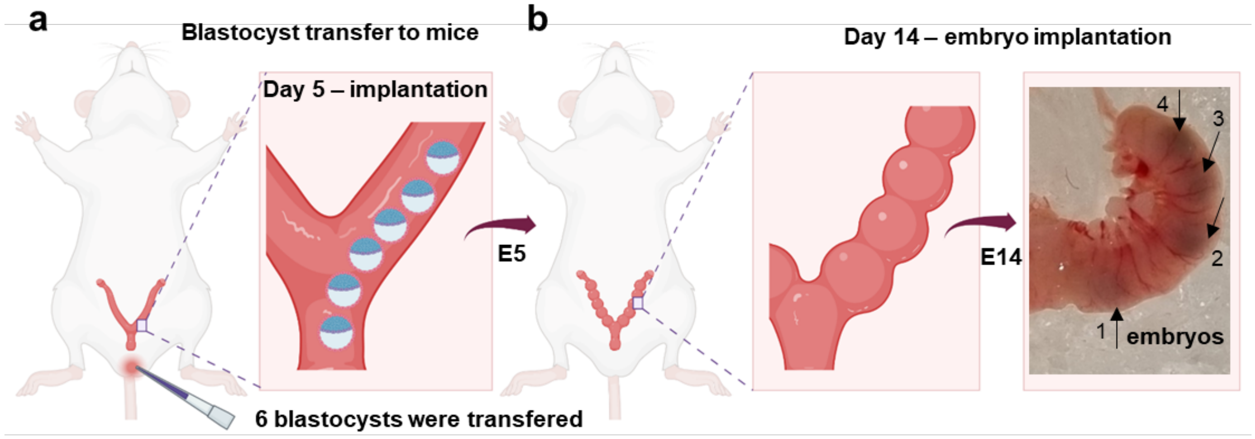


**Fig. S6.** Preliminary tests of embryo transfer and implantation after retrieval of in-spiral cultured embryos. Although development was blocked by day 14, initial signs of successful embryo implantation were observed.

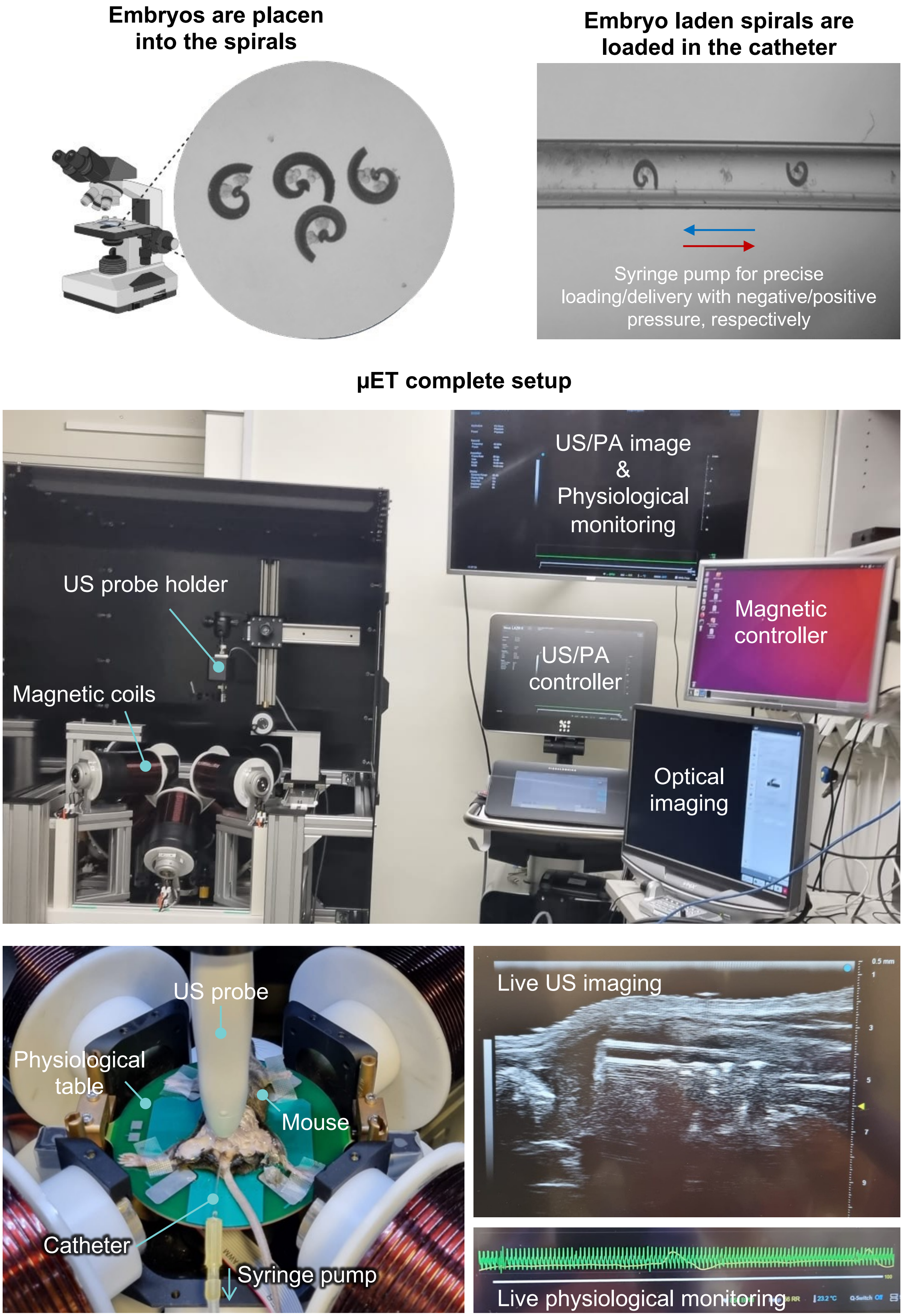
Embryos are placen into the spirals
Embryo laden spirals are loaded in the catheter
Syringe pump for precise loading/delivery with negative/positive pressure, respectively
µET complete setup
US/PA image & Physiological monitoring
US probe holder
Magnetic controller
US/PA controller
Magnetic coils
Optical imaging
US probe
Physiological table
Mouse
Catheter
Syringe pump
Live US imaging
Live physiological monitoring

**Fig. S7.** Setup for in vivo experiments. Electromagnet array surrounding the stage where the mouse is positioned. Various physiological parameters—temperature, respiratory rate, and heart rate—are monitored, while imaging is performed using an ultrasound probe. The animal is anesthetized via oral administration during the procedure. It is also included a representative ultrasound frame showing the uterine horn cavity with two released microrobots, which are actuated using external magnetic fields, while the animal's vital signs are continuously monitored.

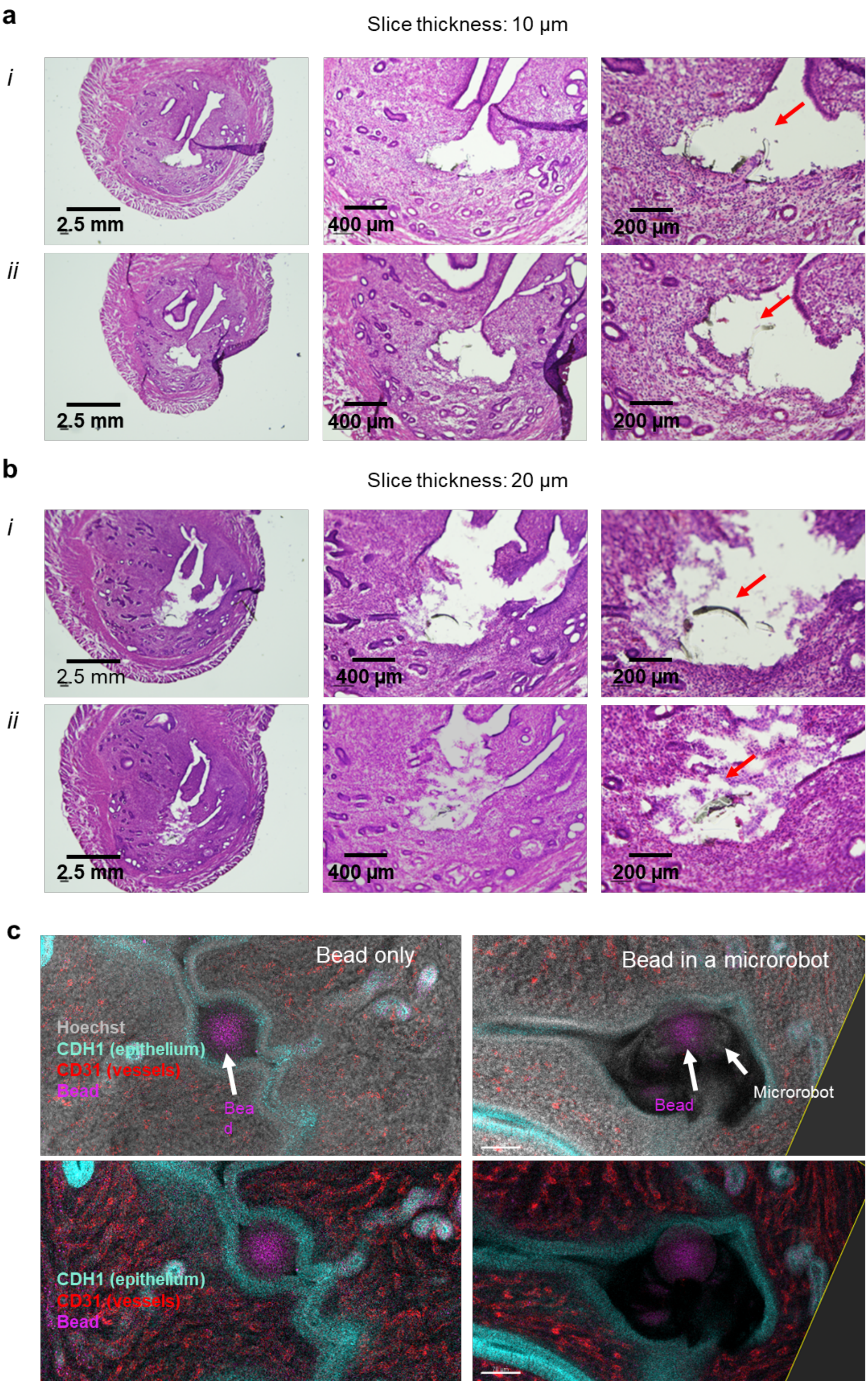
a
Slice thickness: 10 µm
i
ii
2.5 mm
400 µm
200 µm
b
Slice thickness: 20 µm
c
Bead only
Bead in a microrobot
Hoechst
CDH1 (epithelium)
CD31 (vessels)
Bead
Microrobot

**Fig. S8.** Microrobot–tissue interaction studies. (a–b) Histology images (H&E staining) showing cross-sections of the uterine horn, including a section of the spiral microrobot. Tissue slices were prepared at 10 µm (a) and 20 µm (b) thickness before staining. (c) Confocal images using different staining, comparing tissue remodeling around an embryo-mimicking bead alone versus an embryo-mimicking bead loaded inside the spiral microrobot.
